# A serotonergic axon-cilium synapse drives nuclear signaling to maintain chromatin accessibility

**DOI:** 10.1101/2021.09.27.461878

**Authors:** Shu-Hsien Sheu, Srigokul Upadhyayula, Vincent Dupuy, Song Pang, Andrew L. Lemire, Deepika Walpita, H. Amalia Pasolli, Fei Deng, Jinxia Wan, Lihua Wang, Justin Houser, Silvia Sanchez-Martinez, Sebastian E. Brauchi, Sambashiva Banala, Melanie Freeman, C. Shan Xu, Tom Kirchhausen, Harald F. Hess, Luke Lavis, Yu-Long Li, Séverine Chaumont-Dubel, David E. Clapham

## Abstract

Chemical synapses between axons and dendrites mediate much of the brain’s intercellular communication. Here we describe a new kind of synapse – the axo-ciliary synapse - between axons and primary cilia. By employing enhanced focused ion beam – scanning electron microscopy on samples with optimally preserved ultrastructure, we discovered synapses between the serotonergic axons arising from the brainstem, and the primary cilia of hippocampal CA1 pyramidal neurons. Functionally, these cilia are enriched in a ciliary-restricted serotonin receptor, 5-hydroxytryptamine receptor 6 (HTR6), whose mutation is associated with learning and memory defects. Using a newly developed cilia-targeted serotonin sensor, we show that optogenetic stimulation of serotonergic axons results in serotonin release onto cilia. Ciliary HTR6 stimulation activates a non-canonical G_αq/11_-RhoA pathway. Ablation of this pathway results in nuclear actin and chromatin accessibility changes in CA1 pyramidal neurons. Axo-ciliary synapses serve as a distinct mechanism for neuromodulators to program neuron transcription through privileged access to the nuclear compartment.

## Introduction

The primary cilium is a microtubule-based, membrane-bound compartment that extends a few microns from the basal body into the extracellular space (Bornens, 2012). Ciliopathies, genetic disorders caused by mutant proteins related to cilia function, range from embryonic and perinatal death to *situs inversus* (left/right reversed visceral organs), polydactyly, kidney cyst formation, obesity, and neurological deficits such as ataxia and intellectual disability. Many of these phenotypes can be attributed to abnormal embryonic development, since the primary cilia house several key components in the Sonic hedgehog (Shh) pathway that orchestrates the processing and release of cilia-housed GLI1-3 transcription factors affecting growth, division, and differentiation (reviewed in Goetz and Anderson, 2010).

Less is known about the normal function of primary cilia in the mature brain, in which most neurons no longer divide or differentiate. Although cilia are lost in most terminally differentiated adult skeletal and cardiac muscle, they are present in most mature neurons and glia of brain (Guemez-Gamboa et al., 2014). There are a few clues that point to their function in adult brain, but most can be interpreted as late onset phenotypes stemming from developmental abnormalities (e.g., Bardet-Biedl syndrome; Barnett et al., 2002). One important fact is that primary cilia in adult brain are enriched in selected G-protein coupled receptors (GPCRs) for neurotransmitters including dopamine, serotonin, norepinephrine, and somatostatin. Indeed, knock-out of a ciliary-localized GPCR, somatostatin receptor 3 (SSTR3), causes novel object recognition cognitive impairment without grossly affecting brain development (Einstein et al., 2010). To gain insight into the potential functions of neuronal primary cilia in the adult brain, we set out to determine how ciliary signaling events are activated *in vivo* and the sequelae of ligand binding to primary cilia-specific serotonin HTR6 receptors.

## Results

### CA1 neuronal cilia have a preferred orientation

Anti-adenylyl cyclase 3 (ADCY3) antibodies were used to visualize the brain neuronal primary cilia’s distribution and orientation in 200 μm fixed sections (Bishop et al., 2007; Figure 1A). The preferred trajectory of hippocampal pyramidal neuron’s primary cilia is along the basal-apical axis (Figure 1B). This pattern is most striking and uniform in the CA1 region, and least so in the CA2 region. Similar preferred orientations of primary cilia were observed in cortical neurons, which aligned with apical dendrites (Kirschen et al., 2017).

**Figure 1.**
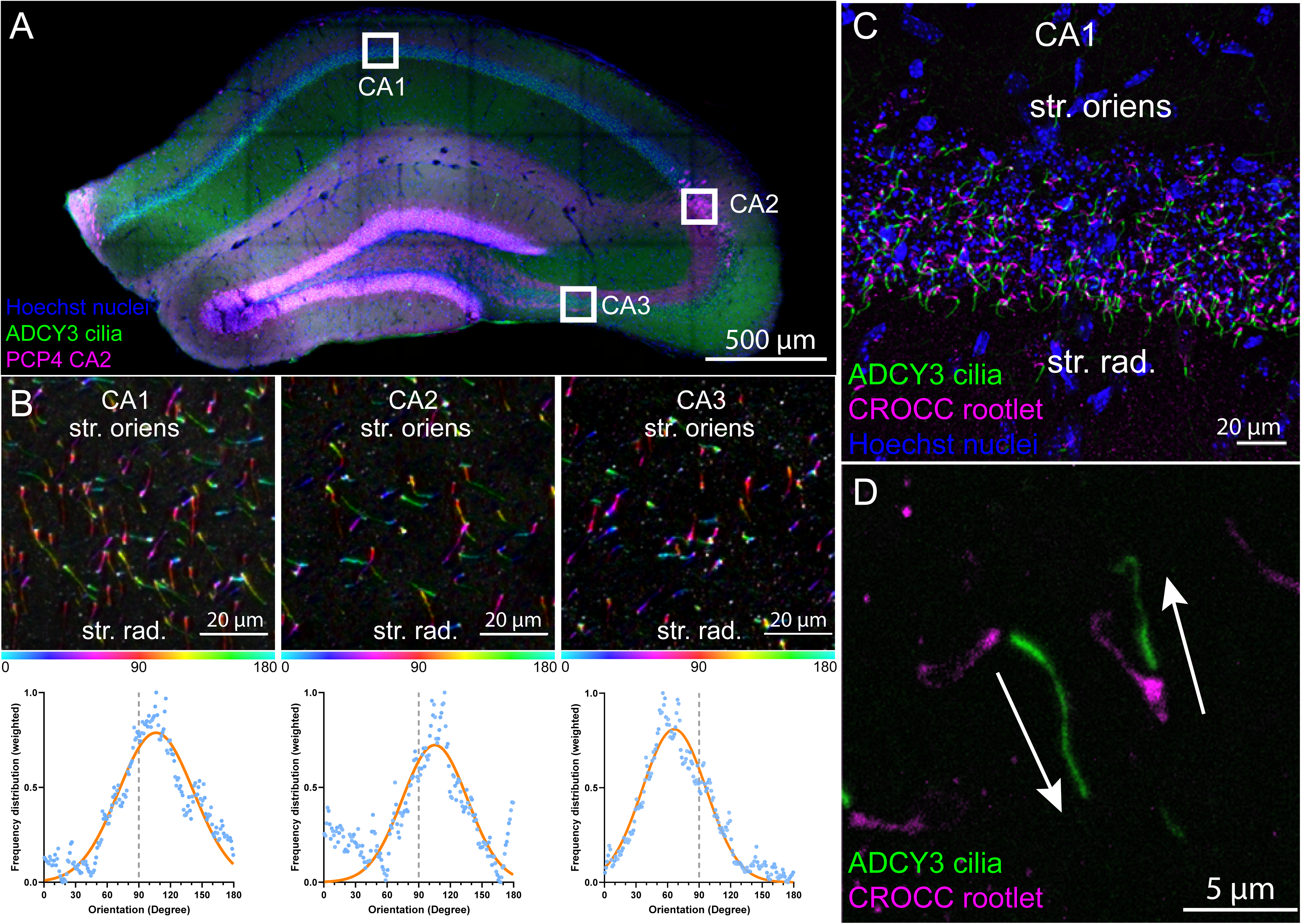
Adult hippocampal pyramidal neuronal cilia are oriented. (**A**) Hippocampal coronal 200 μm-thick section maximum intensity projection (MIP). Green: cilia (ADCY3), red: CA2 (PCP4), blue: nuclei (Hoechst 33342). (**B**) Orientation (structure tensor) analyses of cilia voxels in CA1, CA2, and CA3. 100 μm maximum intensity projection, showing cilia oriented along the basal-apical axis (stratum oriens – stratum radiatum). The images are rotated such that the basal-apical axes are at a 90° angle. Top panel: Color survey of cilia voxels encoded by orientation (hue), coherence (saturation), and fluorescence intensity (brightness). Bottom panel: normalized weighted frequency distribution with basal-apical axes at 90°, showing original data (blue) and fitted gaussian curves (orange). Mean values of the gaussian distributions are 106°, 106°, and 66° for CA1, CA2, and CA3, respectively. The tails of CA2 fit less well than for CA1 and CA3, indicating more heterogeneity in CA2 cilia vectors. (**C**) Labeling of ciliary base. CA1 cilia (green) and Rootletin (magenta; CROCC, ciliary rootlet); nuclei (blue, Hoechst 33342); 50 μm maximum intensity projection. (**D**) Two cilia oriented at 180° in C are magnified. 5 µm maximum intensity projection.

We next asked if CA1 cilia preferentially project from the deep hippocampus (basal side of the pyramidal neurons) to the superficial hippocampus (apical side of the pyramidal neurons). If true, this would suggest a possible morphogen gradient along the basal-apical axis. We labeled the base of the cilia with an antibody against the ciliary rootlet protein, rootletin (CROCC, Figure 1C). Surprisingly, cilia trajectories were largely bidirectional, with about half of cilia projecting to the more superficial layer (stratum radiatum) and the other half projecting to the deeper layer of the hippocampus (stratum oriens, Figure 1D). We hypothesized that cilia trajectory is influenced by special contacts between cilia and nearby structures in the neuropil. To test this hypothesis in mouse brain, we employed volume electron microscopy techniques to visualize neuronal primary cilia and their immediate surroundings.

### FIB-SEM reveals axo-ciliary synapses

We used focused ion beam – scanning electron microscopy (FIB-SEM) to reconstruct the microenvironment of CA1 neuronal primary cilia. In a pilot dataset using 6 nm x 6 nm x 20 nm voxel resolution, we reliably followed two CA1 cilia in a 20 μm x 20 μm x 15 μm volume (Figure 2A). The most tantalizing observation was that axonal varicosities are often close to CA1 pyramidal neuronal cilia. One striking example is shown in Figure 2A: here the two cilia meet at an axonal bouton containing synaptic vesicles and a mitochondrion, reminiscent of classical presynaptic axonal terminals.

**Figure 2.**
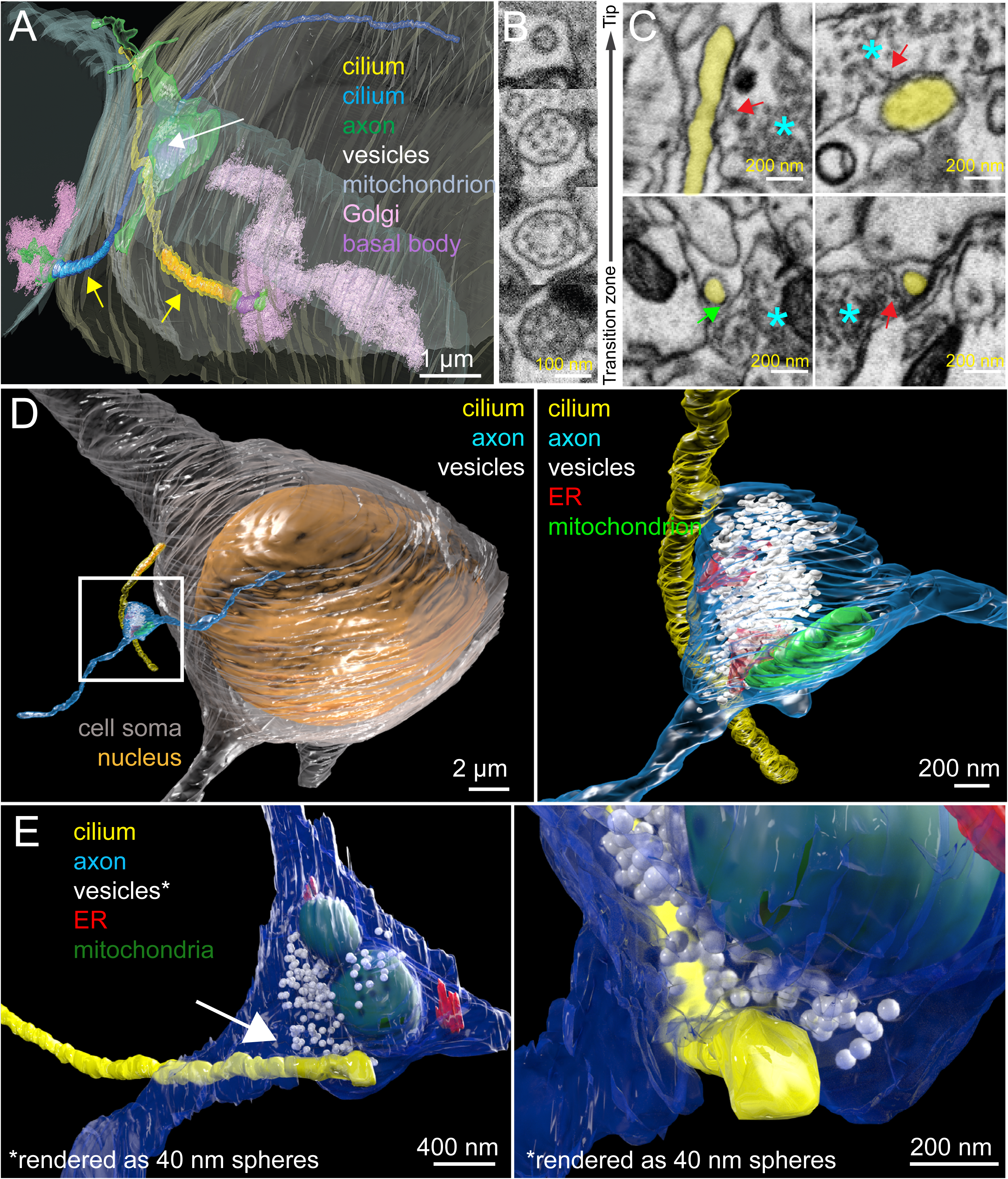
FIB-SEM reveals axo-ciliary synapses. (**A**) Rendering of a 15 μm x 15 μm x 10 μm dataset imaged at 6 nm x 6 nm x 20 nm resolution. Two complete cilia (yellow and blue) arise from the basal bodies, which are surrounded by Golgi-related vesicles and Golgi stacks (pink). Note that the axonal varicosity (green) contains a mitochondrion (lavender) and synaptic vesicles (white, arrow) at the crossing point of the two cilia. Yellow arrows: the portions of the cilia that have identifiable microtubule doublets (2-3 μm; colored in saturated yellow and blue, respectively). (**B**) Primary cilia have a 9+0 microtubule configuration and become 9+1 more distally. No identifiable microtubule doublets are observed in the most distal (6-8 μm) segments (average diameter 100 nm). (**C**) Selected single EM sections of axo-ciliary synapses. Cilium: yellow; axon: cyan asterisk. Top left: An oblique section reconstructed from the volume in (A) to show the longitudinal cross section of the cilium. Note that in some areas, the cilium (yellow) and axonal membrane are in direct contact. Occasional vesicles can be seen within 10-20 nm of the axonal membrane opposing the cilium (red arrow). Bottom left: enhanced contrast at the ciliary membrane next to the axon from volume in A (green arrow). Distance between cilia membrane and axonal membrane at this section is ∼20 nm. Top right and bottom right: selected examples with features suggestive of vesicular docking/fusion at the axonal plasma membrane apposing the cilium from dataset rendered in D-E (red arrow). (**D**) Rendering from an 8 nm x 8 nm x 8 nm isotropic FIB-SEM dataset. An axonal process (cyan) gives rise to a varicosity (white box) that makes direct contact with a pyramidal neuron primary cilium (yellow). White box is magnified in the right panel. Synaptic vesicles in white, endoplasmic reticulum in red, mitochondrion in green. (**E**) Another example of axo-ciliary synapse in the same FIB-SEM dataset. A pyramidal neuronal primary cilium (yellow) originates from the left (base not shown) and makes a contact with an axonal varicosity (blue). The area marked by the white arrow is magnified in the right panel. Synaptic vesicles are rendered as 40 nm spheres to facilitate visualization (white). Note the axonal ensheathment of the cilium, and the proximity of the vesicles to the axonal plasma membrane apposing primary cilia. Endoplasmic reticulum: red, mitochondrion: green.

This raised the question of whether pyramidal neuronal cilia are forming specialized contacts with certain axons, and whether these are specialized sites for neurotransmission. We collected eight FIB-SEM datasets of mature mouse hippocampus at 5.5 nm x 5.5 nm x 15 nm voxel size in 30 μm x 20-30 μm x 20-30 μm volumes. We found that most cilia have contact sites with axonal processes (80%, 25 out of 31). Ciliogenesis of pyramidal neurons start around birth and continue to elongate, until finally shortening at 8-12 weeks (Arellano et al., 2012). In addition, developing brains exhibit more extracellular space than mature brains (Lehmenkühler et al., 1993). We hypothesized that axo-ciliary synapses might be more evident in younger brains. Indeed, axo-ciliary synapses are evident in FIB-SEM images of P14 mouse brain (Figure S1, 83%, 10 of 12 samples). In some cases, pyramidal neuronal cilia and axons appear to travel together (Figure S1A). As in adults, there are mitochondria and ER-PM junctions in axonal processes in contact with primary cilia, resembling classic presynaptic boutons (Figure S1B).

Next, we tested whether fixation artefacts or simply random coincidence were responsible. High-pressure freezing-freeze substitution (HPF-FS) of live samples better preserves ultrastructure and extracellular space (Hoffman et al., 2020; Korogod et al., 2015; Zechmann et al., 2007). However, it is difficult to preserve samples larger than 10 μm without significant formation of ice crystals (Korogod et al., 2015). A hybrid protocol – chemical perfusion followed by HPF-FS provides ultrastructure that is close to direct HPF/FS of live samples for transmission electron microscopy (Sosinsky et al., 2008), but it did not preserve the extracellular space and the contrast was insufficient for FIB-SEM (data not shown). Therefore, we developed a new hybrid protocol to better preserve ultrastructure and extracellular space while providing high contrast for FIB-SEM imaging by employing imidazole and 3-amino,1,2,4 triazole in osmium-based freeze-substitution staining (Figure S2, see Methods). We acquired two isotropic 8 nm x 8 nm x 8 nm FIB-

SEM datasets of adult mouse CA1 samples prepared with the hybrid protocol using enhanced FIB-SEM systems (35 µm x 35 µm x 40 µm and 50 µm x 50 µm x 44 µm). In these two datasets, we identified a total of 27 neuronal primary cilia and similar putative axonal-primary cilia contact sites, or axo-ciliary synapses, in 18 out of 27 neuronal cilia (67%, Figure 2C-E**, Video S1**). These synapses are characterized by a 20-40 nm cleft between the axon and the cilium, flanked by areas in which the axonal membrane and ciliary membrane are immediately in apposition (Figure 2C). In axonal varicosities, synaptic vesicles can be seen within 20 nm of the axonal plasma membrane, or occasionally, appear to be docking or fusing with the plasma membrane, suggestive of vesicular release (Figure 2C, red arrows). Also seen in the axon are endoplasmic reticulum forming junctions with the plasma membrane (ER-PM junctions) and mitochondria, as seen in classical presynaptic terminals (Wu et al., 2017; Figure 2D-E). Interestingly, contrast enhancement is seen at the ciliary membrane next to the axonal varicosity (Figure 2C, arrow). These features resemble classical chemical synapses.

### Axo-ciliary synapses are serotonergic

Since the axons that form axo-ciliary synapses originate from neurons outside the FIB-SEM datasets, we sought to determine the identity of these axons. HTR6, a serotonin G protein-coupled receptor, is highly enriched in neuronal primary cilia (Brodsky and Neumaier, 2017). We hypothesized that the axons in contact with cilia are serotonergic axons that innervate the cilia through the HTR6 receptor. By using an endogenous Htr6-EGFP knock-in mouse line (Nadim et al., 2016), we first confirmed that HTR6 is restricted to pyramidal neuronal primary cilia (Figure 3A). Thirty-five percent (426 out of 1209) of cilia are in close apposition to serotonergic axons (Figure 3B-C), as measured by co-labeling cilia and the serotonin transporter (SERT, SLC6A4, a marker for serotonergic axons) using super-resolution Airyscan confocal microscopy. In addition, all axonal sites opposing cilia contain synaptophysin staining, suggesting that these are serotonin-release sites (Figure 3D-E; Belmer et al., 2017). Since the apposition distance could be subject to variables such as antibody accessibility and optical chromatic aberrations (see Methods), we examined the distribution of the shortest distance to the central axes of serotonergic axons (skeletonized axons) among all ciliary voxels (central axes) and all cilia central axes (skeletonized cilia, Figure 3F-G). Both showed a skewed distribution towards short distances, suggesting that ciliary trajectories are biased towards serotonergic axons. In addition, since ligand stimulation is known to result in ciliary remodeling, including ciliary ectocytosis, decapitation, withdrawal, or shedding (Mirvis et al., 2019; Nager et al., 2017), our analysis of a single snapshot in time might underestimate the frequency of axo-ciliary synapses.

**Figure 3.**
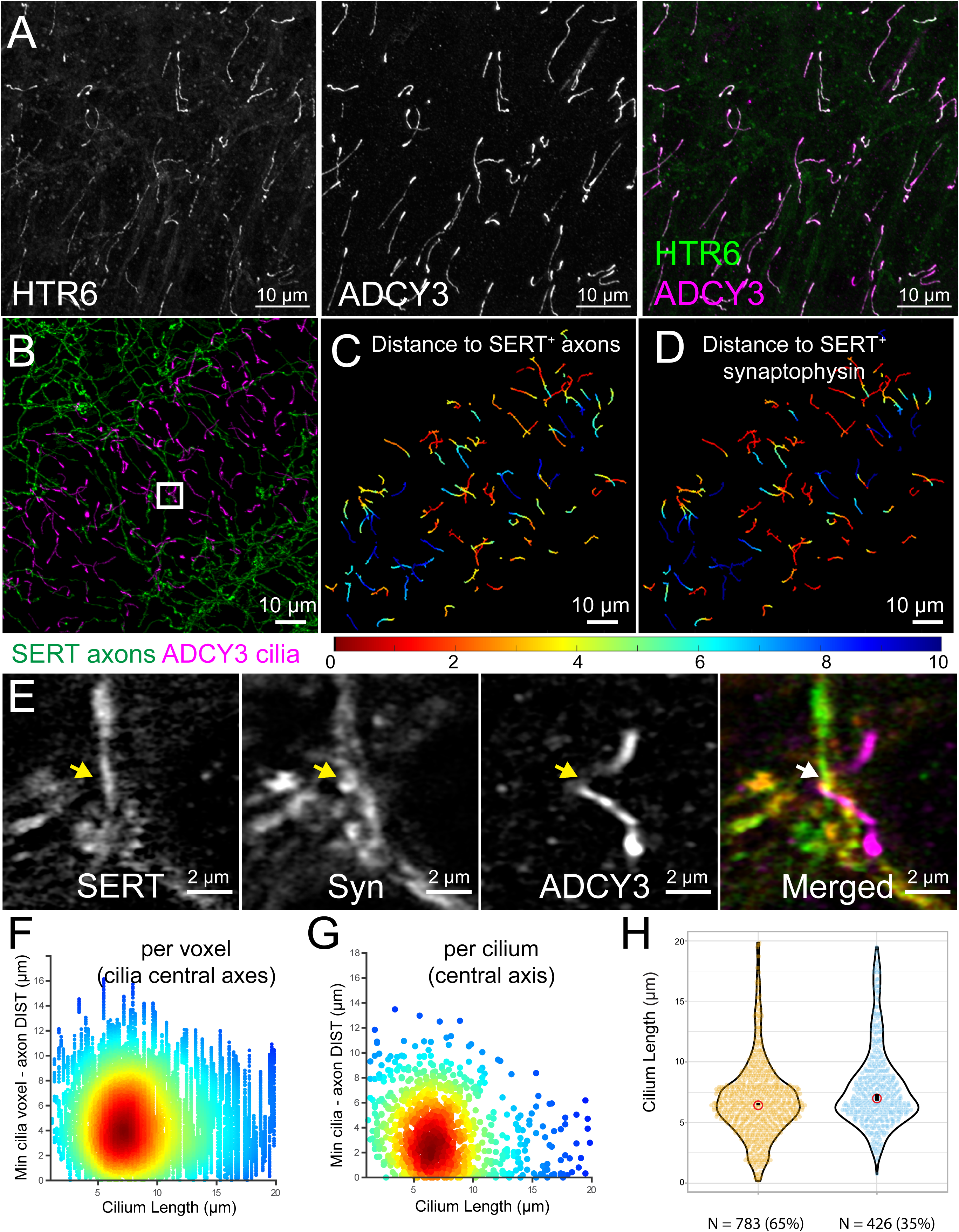
5-hydroxytryptamine (5-HT; serotonin) receptor (HTR6)-expressing primary cilia are in contact with serotonergic axonal varicosities. (**A**) HTR6 (endogenously tagged with EGFP, amplified with anti-GFP antibody and Alexa 488, green in the merged panel) is highly enriched in CA1 neuronal primary cilia (adenylyl cyclase type 3 immunostaining: ADCY3, magenta in the merged panel). (**B**) ADCY3-cilia (magenta) are aligned with serotonergic axons (immunostained with anti-serotonin transporter antibody: SERT, SLC6A4, green). Both neuronal primary cilia and serotonergic axons lie along the basal-apical axis; 20 µm maximum intensity projection. (**C**) cilia in B, color-coded with shortest distance to a serotonergic axon. (**D**) cilia in B, color-coded with shortest distance to a serotonergic axon-associated synaptophysin punctum. (**E**) magnified white box area from B, showing a cilium contacting serotonergic axonal synaptophysin varicosities. Serotonergic axon, synaptophysin, and ADCY3 (cilia) are colored in green, yellow, and magenta in the merged panel, respectively. (**F**) Density plot showing the relationship between cilium length and the shortest distance to a serotonergic axon on a per voxel basis (cilium central axis). (**G**) Density plot showing the relationship between cilia length and shortest distance to a serotonergic axon per cilium (central axis). Notice the lack of a linear correlation, and the skewed distribution towards shorter distances. (**H**) Violin plots showing the distribution of ciliary length of serotonergic axon-contacting and non-contacting cilia. Red circle: median; black bar: 95% confidence interval of the median. The difference in the shapes of the violin plots and the length of the 95% bars reflect the greater variance of contacting cilia.

We noticed that the axon-contacting cilia are slightly longer than the non-contacting cilia (Figure 3H; median length 7.0 versus 6.4 μm, p<0.0001, two-tailed Mann-Whitney U test). However, this length difference of <1 μm is unlikely to explain why some cilia are in contact with serotonergic axons while others are not. Indeed, we failed to detect any significant correlation between cilia length and the shortest distance to serotonergic axons (Figure 3G, Pearson correlation coefficient r = -0.19), suggesting that simply increasing ciliary length would not result in an increased likelihood of contacts with serotonergic axons. Interestingly, contacting cilia exhibit greater variance in length (standard deviation: 3.3 μm for contacting cilia, versus 3.0 μm for non-contacting cilia; two-group distribution comparison p-value with permutation test <0.0001), this implies that serotonergic axon-contacting cilia may receive higher levels of stimulation and thus have more frequent changes in length. Altogether, these results suggest that CA1 pyramidal neuronal cilia receive serotonergic innervation from the raphe nuclei (Muzerelle et al., 2016).

### Activation of serotonergic axons releases serotonin onto cilia

The ultrastructural and super-resolution analyses above provide anatomical evidence of axo-ciliary synapses. Next, we sought to determine if serotonin is released onto cilia upon the activation of serotonergic axons. Fluorescent reporters often do not traffic to cilia (Delling et al., 2013), so we first fused an extracellular serotonin sensor (Unger et al., 2020) onto the N-terminus of HTR6. Unfortunately, the HTR6 receptor was aberrantly targeted. As an alternative, we engineered a ciliary-targeted serotonin sensor based on the GPCR-activation-based (GRAB) strategy with the HTR6 receptor as the scaffold (GRAB-HTR6-PM; Wan et al., 2021). We first expressed this sensor in HEK-293T cells, which trafficked well to membranes with an ∼150% fluorescence increase in response to saturating 5-HT (EC_50_ = 84 nM; Figure S3; human HTR6 receptor K_D_ = 37 nM; Monsma et al., 1993). Therefore, we reasoned that this sensor could detect ciliary serotonin changes at physiologically relevant levels. We then removed the IgK leader sequence and added a HaloTag on the C-terminus to better visualize cilia with bright Janelia Fluor dyes (Grimm et al., 2015; Zheng et al., 2019). This resulted in robust cilia targeting in RPE-1 cells and neurons, as when HTR6 is expressed (Figure 4A). The EC50 for the cilia-targeted HTR6-GRAB-cilia sensor is 28 nM, with up to 40% fluorescence increase per cilium in response to saturating doses of 5-HT using 3D Airyscan (Figure 4B). The smaller increase in maximum fluorescence of the cilia-targeted sensor (40% versus 150%) may be attributed to a different lipid composition of the ciliary membrane (reviewed in Conduit and Vanhaesebroeck, 2020) and/or the increased bleaching due to laser scanning/Z-stack imaging required to 3-dimensionally image single cilia.

**Figure 4.**
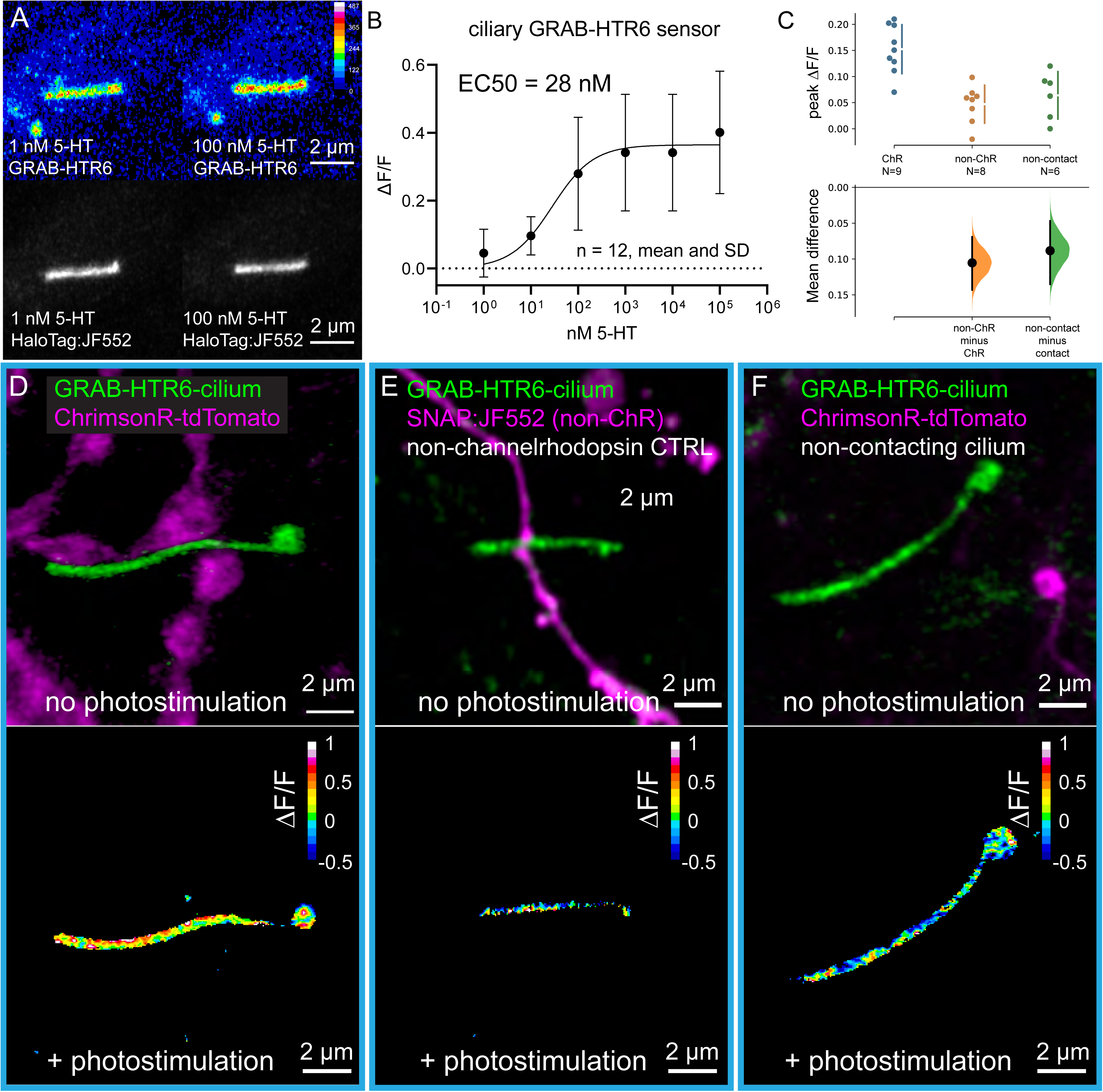
Activation of serotonergic axons releases serotonin onto cilia. (**A**) RPE-1 cells stably expressing a Tet-inducible HTR6-GRAB-cilia-HaloTag serotonin sensor. 100 nM application results in increased GFP fluorescence. HaloTag:JF552 is used to reliably identify and segment cilia. (**B**) Titration curve of the sensor. (**C**) Statistical analyses of ciliary serotonin levels with optogenetic stimulation of serotonergic axons, shown in the Cumming estimation plot. The raw data is plotted on the upper axes. On the lower axes, mean differences are plotted as bootstrap sampling distributions. Each mean difference is depicted as a dot. Each 95% confidence interval is indicated by the ends of the vertical error bars. Peak ΔF/F is calculated by filtering the averaged measurements using a 1 μm circle at the contact sites (ChrimsonR and non-ChrimsonR contacting cilia controls), or the location on the cilium that is closest to a serotonergic axon (ChrimsonR non-contacting cilia) using a low pass filter (see Figure S5 and Methods). (**D-F**) Top: A cilium expressing the HTR6-GRAB-cilia sensor in contact with ChrimsonR-tdTomato-expressing serotonergic axons (**D**), in contact with a SNAP:JF552 labeled serotonergic axon (non-channelrhodopsin control, **E**), and a cilium distant from a ChrimsonR-tdTomato-expressing serotonergic axon (**F**). Bottom: color maps showing ΔF/F during optogenetics stimulation (after 25 pulses at 1Hz), corresponding to the 3 different examples in the top row, respectively.

We attempted imaging ciliary serotonin dynamics in acute hippocampal slices with mice injected with adeno-associated virus (AAV) carrying the sensor construct. However, we could not achieve good signal-to-noise ratios in deeper areas (>20 μm), in which surface damage can be avoided with either Airyscan or lattice light-sheet microscope with adaptive optics (data not shown). Therefore, we developed an in-vitro system of serotonergic axo-ciliary synapses by co-culturing hippocampal and serotonergic neurons from the midbrain (Figure S4). We expressed the cilia-serotonin sensor GRAB-HTR6-cilia and ChrimsonR, a red-shifted channelrhodopsin (Klapoetke et al., 2014), in hippocampal and serotonergic neurons, respectively. Using Airyscan, we detected serotonin release onto cilia that were in synaptic contact with serotonergic axons (average peak ΔF/F = 0.15, Figure 4C-D; Figure S5). This increase is diminished in non-channelrhodopsin controls (mean difference, or effect size by estimation statistics = -0.10, 95% CI = -0.14 to -0.7, permutation test p-value = 0.0002, two-tailed Mann-Whitney test p-value = 0.0009, Figure 4C,E; Figure S5). Interestingly, the serotonin release is attenuated on cilia distant from serotonergic axons (mean difference, or effect size by estimation statistics = -0.09, 95% CI = -0.14 to -0.05, permutation test p-value = 0.005, two-tailed Mann-Whitney test p-value = 0.008, Figure 4C,F; Figure S5). Although the differences are near the limit of detection due to the sensitivity of the sensors/detectors and the small volumes being measured, the data suggest that firing of serotonergic axons results in serotonin release on hippocampal neuronal cilia. Future experiments may be possible using 5HT3 channel-containing sniffer pipettes placed in apposition to the releasing membrane to detect serotonin, although surface accessibility currently limits such approaches.

### Serotonin stimulation activates a neuronal ciliary HTR6-G**_αq/11_**-Trio-RhoA pathway

To examine the functional significance of the axo-ciliary synapses, we studied serotonin-induced HTR6 activation on cilia. HTR6 is characterized as a G_αs_-coupled GPCR, activating adenylyl cyclase to increase cAMP when over-expressed on the plasma membrane of dividing HEK cells (Boess et al., 1997). However, HTR6 activation does not increase cAMP when it is localized in cultured cell primary cilia (Jiang et al., 2019). However, GPCRs may interact with multiple G-proteins (Flock et al., 2017; Masuho et al., 2015; Okashah et al., 2019). Indeed, GPCR-G protein coupling differs in some ciliated *vs* non-ciliated cells (Masyuk et al., 2013) and the same GPCR might be coupled to different Gα-subunits (Hilgendorf et al., 2019) on the plasma membrane *vs* the cilia. Therefore, we hypothesized that cilia-localized HTR6 interacts with a different G_α_ subunit. The G_α11_-subunit (GNA11) was previously identified as a binding partner of the endogenous HTR6 in the brain through affinity purification and mass spectrometry (Nadim et al., 2016), contrasting with AP-MS data obtained by heterologous expression of HTR6 in dividing HEK cells, which identified the G_αs_-subunit (GNAS), as a binding partner (Meffre et al., 2012).

G_αq/11_ can also activate the Trio-RhoA pathway in *C. elegans* and in G_αq/11_-constitutively active mutant uveal melanoma cells, as determined through a forward genetic screen and a genome-wide siRNA screen, respectively (Feng et al., 2014; Williams et al., 2007). Trio was identified in the HTR6-ciliome (Kohli et al., 2017) and is detected in cilia by immunofluorescence in HTR6-HaloTag RPE-1 cells and in WT cultured hippocampal neurons (Figure S6A). Indeed, serotonin activates RhoA in HTR6-overexpressing HEK cells and in neurons in which the receptor is distributed throughout the plasma membrane and neuronal processes, suggesting that HTR6 may signal through the G_αq/11_-Trio-RhoA pathway (Rahman et al., 2017). We thus hypothesized that neuronal ciliary HTR6 expressed at the endogenous levels might activate RhoA through G_αq/11_-Trio.

To our knowledge, RhoA activity has not been directly measured in primary cilia. We first tested several translocation based RhoA sensors with fluorescent proteins fused to a protein domain that binds to active RhoA (Mahlandt et al., 2021). However, these sensors caused either significant cell death or formed cytoplasmic aggregates in neurons (data not shown). We then tested FRET-based RhoA sensors, in which a pair of fluorescent proteins is fused to RhoA and a protein domain that binds to active RhoA, respectively. We targeted a FRET-based RhoA sensor (Bindels et al., 2017) to the cilia by fusing it to HTR6 and expressed it in RPE-1 cells (Figure S6C). To better recapitulate the serotonin release at the axo-ciliary synapses, we considered employing caged serotonin next to the cilium using UV-releasable (N)-1-(2-nitrophenyl) ethyl (NPEC)-caged serotonin (Breitinger et al., 2000). To avoid damaging UV radiation, we synthesized a new photoactivatable, caged serotonin molecule that can be cleaved by 405 nm laser light (Figure S6B). Caged serotonin stimulation (0.5 Hz) immediately adjacent to the cilium elicited a pulsatile increase in RhoA activity, returning to near baseline upon cessation (Figure S6C-E). However, uncaging suffered from a high failure rate and the average FRET ratios across cilium can be significantly affected by just a few pixels due to the small size of cilium.

We sought to obtain better measurements of ciliary RhoA by fluorescence-lifetime measurements (FLIM/FRET) to minimize the effect from donor bleaching and better account for the difference in sensor levels. In this measurement, we expect the donor lifetime to decrease from FRET. As cilia often span multiple Z-levels while FLIM is normally carried out without optical sectioning, we first tested whether FLIM with optical sectioning across the Z axis can be achieved by using a fast FLIM system equipped with a pulsed white light laser (Rolf et al., 2021). We were able to reconstruct whole HEK293A cells with HTR6-RhoA sensor expression through FLIM imaging (Figure 5A). The Arl13b-RhoA sensor is functional in cilia since stimulation by a RhoA activator (Flatau et al., 1997; Schmidt et al., 1997) decreased the fluorescence lifetime of the donor significantly (the effect size, or mean difference by estimation statistics = -60ps, 95% CI = -100ps to -24ps, permutation test p-value = 0, two tailed Mann-Whitney test p-value = 0.002, Figure 5B-C). HTR6-RhoA cilia have higher RhoA activity than Arl13b-RhoA cilia, suggesting that over-expression of HTR6 results in constitutive activity (mean difference, or effect size by estimation statistics = -125ps, 95% CI = -162ps to -93ps, permutation test p-value = 0, Mann-Whitney test p-value <0.00001, Figure 5D-E), as commonly seen in GPCR signaling (reviewed in Seifert and Wenzel-Seifert, 2002).

**Figure. 5.**
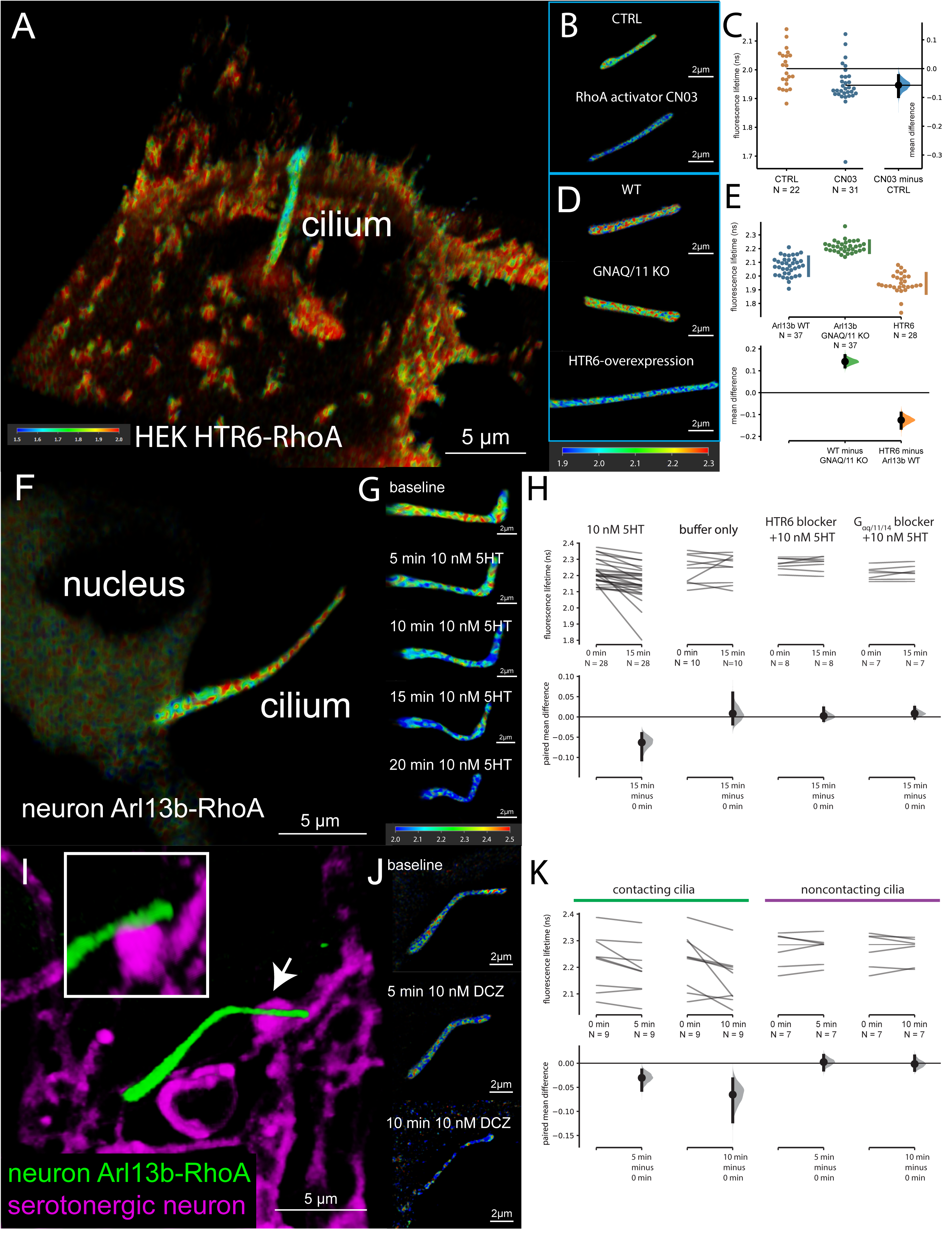
Serotonin stimulation of ciliary HTR6 activates RhoA in cilia. (**A**) HEK293A cells stably expressing the HTR6-RhoA FRET/FLIM sensor. A single cilium arises from the dome-shaped cell soma (FLIM). (**B-C**) Cilia-targeted Arl13b-RhoA sensor responds to RhoA activation. The mean difference between control and CN03-(RhoA activator) treated cells is shown in the Gardner-Altman estimation plot. Both groups are plotted on the left axes; the mean difference is plotted on a floating axis on the right as a bootstrap sampling distribution. The mean difference is depicted as a dot; the 95% confidence interval is indicated by the ends of the vertical error bar. (**D-E**) GNAQ/11 KO and HTR6-overexpression decreases and increases RhoA activity, respectively. The mean differences for 2 comparisons against the shared control Arl13b WT are shown in the Cumming estimation plot. The raw data is plotted on the upper axes. On the lower axes, mean differences are plotted as bootstrap sampling distributions. Each mean difference is depicted as a dot. Each 95% confidence interval is indicated by the ends of the vertical error bars. (**F-H**) 10 nM 5HT stimulation of neuronal cilia increases ciliary RhoA activity. This effect is blocked by either HTR6 blocker SB258585 (100 nM) or the G_αq/11_ blocker YM-254890 (1 µM). (**I-K**) Chemogenetic stimulation of serotonergic axons increases ciliary RhoA activity in contacting cilia but not in non-contacting cilia. Arrow in I points to area magnified in the inset, which is shown at an oblique angle to demonstrate the close apposition of axon and cilium at the synapse. Contrast is enhanced in the 10 min time point in J. For H and K, the paired mean differences for comparisons are shown in the Cumming estimation plot. The raw data is plotted on the upper axes; each paired set of observations is connected by a line. On the lower axes, each paired mean difference is plotted as a bootstrap sampling distribution. Mean differences are depicted as dots; 95% confidence intervals are indicated by the ends of the vertical error bars.

We then measured serotonin-dependent RhoA activity in neuronal cilia. We expressed the Arl13b-RhoA sensor, as cultured hippocampal neurons have ciliary HTR6 (Figure 5F). Due to instrument limitations, we were unable to perform photo-uncaging concurrently with FLIM measurements. Instead, we used a low concentration of 5HT (10 nM; rat receptor K_D_ = 12.7 nM; Boess et al., 1997) to minimize receptor desensitization and better recapitulate the pulsed nature of serotonin release by axonal firing. Ten nanomolar 5HT stimulation reliably increased RhoA activity in neuronal cilia in 5 to 15 minutes (the effect size, or the estimation statistics = -63ps, 95% CI = -106ps to -41ps, permutation test p-value = 0, Wilcoxon p-value = 0.00001, Figure 5G-H). Adding a HTR6 blocker SB258585 (100 nM, Hirst et al., 2000) 5 min before 10 nM 5HT application largely abolished this effect (Figure 5H), suggesting that the increase in RhoA requires HTR6. We then asked if the increase required G_αq/11_. Pretreatment with a G_αq/11_ inhibitor, YM-254890 (1 μM; Nishimura et al., 2010; Takasaki et al., 2004) mitigated the RhoA increase in neuronal cilia (Figure 5H). G_αq/11_ KO HEK293A cells have significantly lower ciliary RhoA activity; Figure 5D-E). Lastly, pre-treatment with YM-254890 abolished RhoA spikes seen in RPE-1 cells (Figure S6D-E). Together, these data suggest that serotonin stimulation results in G_αq/11_ -dependent RhoA activation in cilia.

Next, we tested ciliary RhoA activity upon chemogenetic activation of serotonergic axons in the hippocampal neuron – raphe neuron co-culture system. We expressed the Arl13b-RhoA sensor and an excitatory Designer Receptors Exclusively Activated by Designer Drugs (DREADD) hM3Dq (Armbruster et al., 2007) in hippocampal neurons and raphe neurons, respectively. Application of 10 nM hM3Dq DREADD agonist deschloroclozapine (DCZ, Nagai et al., 2020) increased RhoA activity in cilia that apposed serotonergic axons within 5 min (Figure 5I-K). In some cases, we observed ciliary withdrawal and/or retraction, or receptor retrieval from the cilia. In contrast, there was no detectable increase in RhoA activity in non-contacting cilia (Figure 5K). This suggests that the ciliary RhoA activation is under spatial and temporal control of the activity of serotonergic axons.

### Nuclear actin-associated changes and decreased chromatin accessibility after ciliary HTR6 ablation

RhoA activation is known to induce cytoplasmic F-actin polymerization and myosin activation (reviewed in Hodge and Ridley, 2016). As cilia originate from the neuronal cell somata, we hypothesized that ciliary RhoA signaling can affect somatic actin-related structures. The most notable examples of actin-related structures in neuronal cell somata are actin lattices revealed by dSTORM imaging (Han et al., 2017). These lattices were revealed using antibodies against the actin binding protein, adducin, and resemble the classic lattices seen in red blood cells (Bennett and Gilligan, 1993; Pan et al., 2018). RhoA activation can phosphorylate adducin through Rho-associated kinase and increase its affinity towards F-actin (Fukata et al., 1999). Consistent with Han et al. (2017), we detected adducin plasma membrane labeling in neuronal cell somata (Figure 6A). When we treated cultured hippocampal neurons with the HTR6 antagonist SB-742457 (Upton et al., 2008), however, we did not see significant changes in plasma membrane adducin, but nuclear adducin was enriched in a small subset of neurons (Figure 6B). This change is reminiscent of nuclear translocation of adducin reported in cultured cells that affect cell growth and division (Chen et al., 2011; Liu et al., 2017).

**Figure 6.**
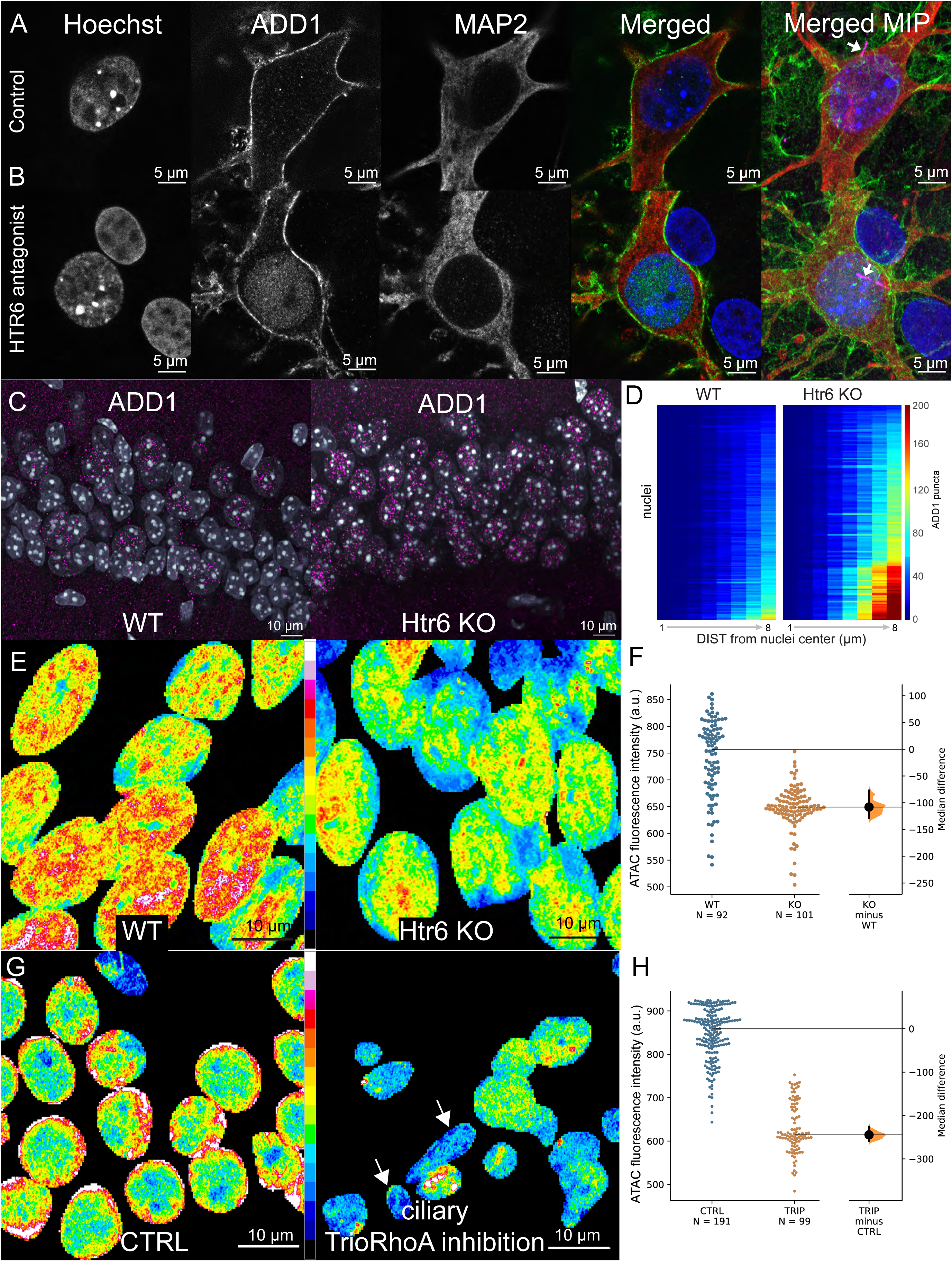
Modulation of the HTR6 signaling axis alters adducin localization and chromatin accessibility. (**A-B**) DIV28 hippocampal neurons were treated either with 0.01% DMSO control (A), or 100 nM SB-742457 (B) for 20 min. In DMSO-treated neurons, adducin is primarily at the plasma membrane. In contrast, in some SB-742457-treated neurons, there is significant nuclear labeling. Pyramidal neurons are identified by MAP2 labeling in both cases. Color scheme in merged panels: Blue: Hoechst 33342, green: adducin, red: MAP2. Cilia are colored in magenta (ADCY3 staining, arrow) in the merged maximum intensity projection (MIP) of the entire neuron, while other panels are single optical sections through the middle of the nucleus. (**C**) Htr6 KO mice exhibit increased numbers of pyramidal neurons with nuclear adducin (ADD1) puncta. (**D**) Heatmap of number of ADD1 nuclear puncta (color-coded) using the center of nucleus (represented in rows) with increasing distance from 1 to 8 μm (represented in columns). There is increased density of puncta in KO cells. (**E**) Representative single optical Airyscan section of CA1 pyramidal neurons showing ATAC-see labeling with ATTO-590 dye (segmented with a nuclear mask, see Methods). The labeling is significantly decreased in the KO mice, which is quantified in (**F**). (**G**) Representative single optical Airyscan section of CA1 pyramidal neurons showing ATAC-see labeling with ATTO-565 dye (segmented with a nuclear mask, see Methods). The labeling is significantly decreased in the mice fed with doxycycline-containing food to induce the expression of the cilia-targeted TrioRhoGEF inhibitor. Notice the decreased cell density and small, pyknotic nuclei (arrow). The difference between nuclei that are > 8 μm in diameter (not judged pyknotic) is quantified in the estimation statistics plot (**H**). For F and H, the median difference between the two groups is shown in the Gardner-Altman estimation plot. Both groups are plotted on the left axes; the median difference is plotted on a floating axis on the right as a bootstrap sampling distribution. The median difference is depicted as a dot; the 95% confidence interval is indicated by the ends of the vertical error bar.

As cells grown on hard surfaces such as glass can alter actin dynamics, we examined adducin staining patterns in the native hippocampal environment. Surprisingly, we did not detect plasma membrane adducin staining in the neuronal cell somata in the hippocampus, but pyramidal neurons exhibit variable numbers of clear nuclear adducin puncta (Figure 6C). In Htr6 knockout (KO) mice pyramidal neurons, the density of adducin puncta increased significantly (Figure 6D, effect size, or mean difference by estimation statistics within 5 μm radius from the center of nuclei = 25.5, 95% CI = 20.4 to 47.6, permutation test p-value = 0, two-tailed Mann-Whitney test p-value<0.0001), consistent with our HTR6 antagonist experiments. This suggests that ciliary HTR6 signaling is linked to nuclear actin.

Alterations in nuclear actin modify global chromatin (Plessner and Grosse, 2019; Zhao et al., 1998). Increased nuclear actin by disruption of cytoplasmic actin can alter cellular state to drive differentiation (Sen et al., 2017), while reducing nuclear actin can result in cell quiescence (Spencer et al., 2010). To test chromatin remodeling directly in the brain, we employed ATAC-see, a technique that utilizes a hyperactive transposase mutant with fluorescently labeled oligonucleotides (Chen et al., 2016; Xie et al., 2020). The transposase has high reactivity only to open chromatin regions, resulting in fluorescent labeling of accessible chromatin areas in cells. We performed ATAC-see with super-resolution Airyscan on age- and gender-matched samples from WT and Htr6 KO adult mice. In Htr6 KO CA1 samples, the labeling intensity was significant less than the intensity in WT samples (the effect size, or median difference by estimation statistics = -108, 95% CI = -128 to -77, permutation test p-value = 0, two tailed Mann-Whitney test p-value < 0.0001, Figure 6E-F). This change in ATAC-see labeling is reminiscent of the difference between early and late phase G1 cells, in which cells decondense their chromatin after mitosis (Chen et al., 2016). We conclude that ablation of the HTR6 receptor results in altered patterns of accessible chromatin.

To further test if chromatin changes were indeed ciliary RhoA-dependent, we expressed a cilia targeted TrioRhoGEF inhibitor peptide (Bouquier et al., 2009) under a tetracycline inducible promoter. In adult mice fed for 1-week with doxycycline containing food, there was notable neuronal loss with pyknotic nuclei (Figure 6G). The remaining, comparatively healthier neurons had decreased ATAC-see labeling intensity compared to non-doxycycline food treated mice (the effect size, or median difference by estimation statistics = -244, 95% CI = -259 to -225, permutation test p-value = 0, two tailed Mann-Whitney test p-value < 0.0001, Figure 6G-H). Taking all our results together, the most parsimonious conclusion is that primary ciliary HTR6 signaling controls chromatin remodeling states via a G_αq/11_-Trio-RhoA pathway.

### Htr6 KO is associated with loss of accessibility of cis-regulatory elements of genes related to learning and memory

To further characterize the chromatin accessibility changes in Htr6 KO mice, we utilized single-cell ATAC sequencing. We performed nuclei isolation, tagmentation, and sequencing in WT and Htr6 KO hippocampus, followed by pooled data processing (Figure 7A-B). We observed 19 unique cell clusters in pooled WT and KO nuclei (Figure 7C). Cell types were assigned based on differentially enriched marker peaks and known gene expression profiles of the mouse hippocampus (Figure S7 and **Table S1**, see Methods; Zeisel et al, 2018). CA1 neurons were identified as clusters 1, 9, and 13 using GPR161 and Ndst3 as marker genes. Interestingly, the difference between WT and Htr6 KO CA1 neurons are readily apparent in pooled clustering. While cluster 9 is only in WT neurons (n=622), cluster 1 is mostly present in the KO sample (n=1652 and n=117 in KO and WT, respectively, Figure 7D). We compared accessible loci between cluster 1 (KO) and 9 (WT), between all WT CA1 and KO CA1 neurons, and between WT and KO cluster 13. Consistent with our data from ATAC-see, there is significant net loss of accessible loci in all three comparisons in KO neurons (Figure 7E). For all CA1 neurons, KO cells have 3340 loci with lower accessibility than WT, while having 1111 loci with higher accessibility levels.

**Figure 7.**
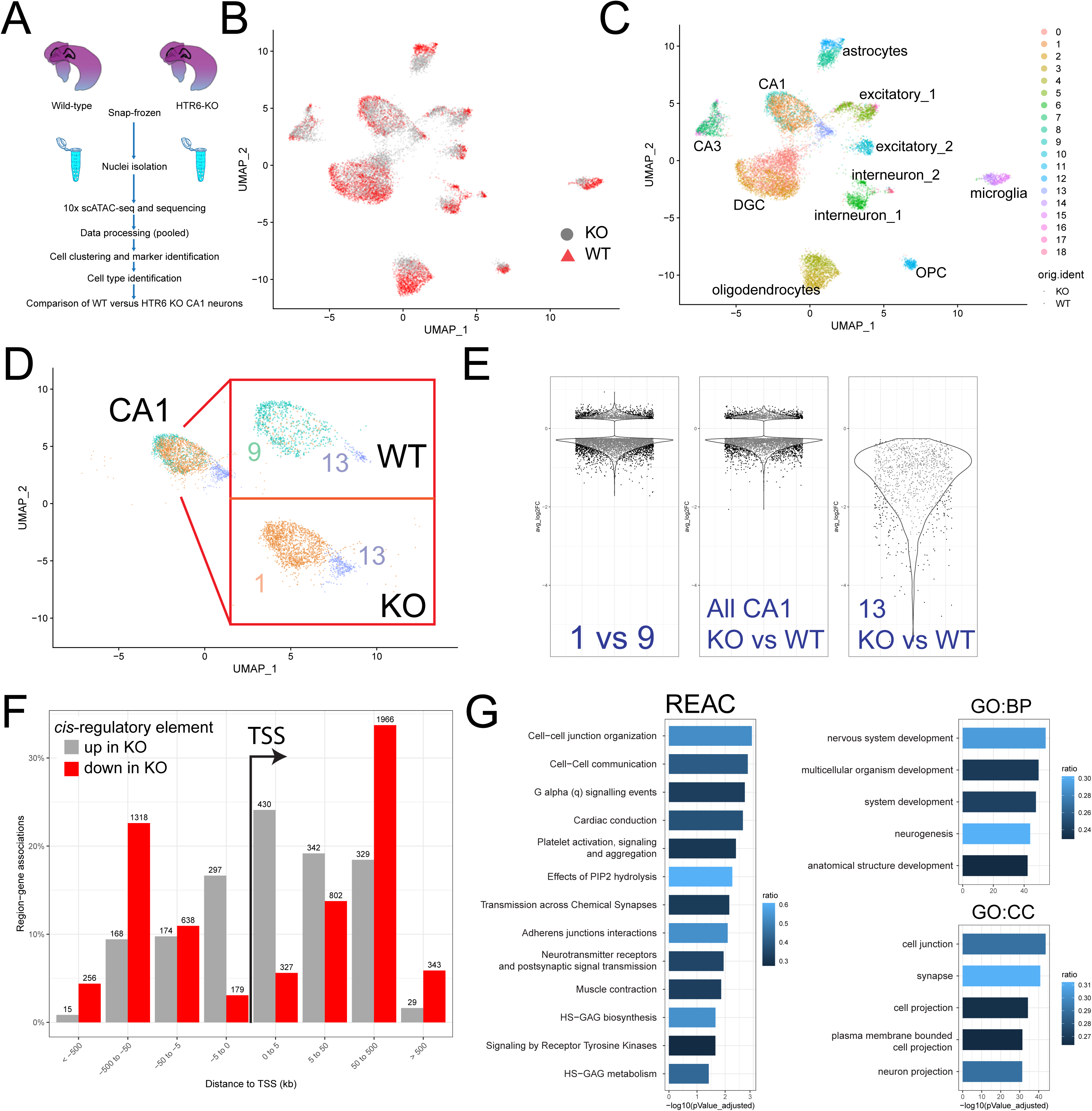
Single-cell ATAC sequencing reveals loss of accessibility to long-range enhancer elements and activity changes in genes associated with synapses and the perineuronal net. (**A**) Overall workflow. (**B**) Pooled clustering among all nuclei (KO and WT). (**C**) All 19 detected clusters were merged into defined hippocampal cell types based on marker peaks (see Methods). (**D**) WT CA1 (cluster 9) and KO CA1 (cluster 1) are enriched in different clusters (pooled). The number of nuclei for cluster 13 increased in KO. (**E**) Analyses of differential peaks between KO and WT, showing on average a 3-fold greater number of less accessible *vs* more accessible loci. (**F**) Less accessible loci in KO are mostly distant from transcription start sites (TSS; red), while more accessible loci are primarily near promoters (gray). (**G**) Gene ontology and Reactome terms enriched in genes affected by the lost loci. REAC: Reactome; GO: Gene Ontology; BP: biological process.; CC: cellular components.

Most of the less accessible loci are distant to transcription start sites (TSS), which may be long range enhancer elements (40% being > 50kb upstream of TSS, 27% being > 50 kb downstream of TSS, with only 9% within ±5kb of TSS sites). In contrast, 40% of more accessible loci are within ±5kb of TSS, or around the promoter area (Figure 7F). This pattern is strikingly similar to that found between low and hi-G1 cells (Chen et al, 2016). The pattern suggests that the ciliary HTR6 signaling axis in mature neurons might impact transcription primarily through long-range enhancers, such as the prototypic *Shh* enhancer that controls the expression of *Shh* gene during development (reviewed in Schoenfelder and Fraser, 2019), Gli repressors through altering enhancer activities (Li et al., 2016), or ciliary FFAR4 receptor-driven adipogenesis through CTCF-binding enhancers (Hilgendorf et al., 2019).

Finally, we sought to determine genes that are impacted by loss of Htr6. By using nearby genes of the less accessible loci (n = 5829, GREAT analysis, McLean et al., 2010), we performed Gene Ontology (Ashburner et al., 2000; The Gene Ontology Consortium et al., 2020) and Reactome (Jassal et al., 2019) term enrichment analyses using g:Profiler (Raudvere et al., 2019). The significantly enriched terms include cell-cell junction and communication, G_αq_ signaling pathways, heparan sulfate and heparin (HS-GAG) biosynthesis and metabolism, cell projection, and synapses (Figure 7G). Notably, a recent study showed that depleting cilia completely in CA1 neurons through IFT88 shRNA knock-down results in defective extracellular matrix/perineuronal net and memory impairment (Jovasevic et al., 2021), suggesting a potential convergent path for these different perturbations. Together, these changes corroborate the synaptic and cognitive deficits reported in Htr6 KO mice (Sun et al., 2021).

## Discussion

We presented evidence of synapses between raphe serotonin-secreting axonal neurons and HTR6 serotonin receptor-expressing primary cilia of CA1 pyramidal neurons. We 1) identified axo-ciliary synapse structures; 2) identified primary cilia HTR6 receptors adjacent to serotonergic axonal varicosities containing vesicles and other markers of synapses; and 3) demonstrated serotonin release onto cilia upon optogenetic stimulation of serotonergic axons. We provided evidence that ciliary HTR6 can activate the non-canonical G_αq/11_-Trio-RhoA pathway in primary cilia. Abrogation of this pathway in mature neurons alters nuclear actin via adducin translocation, thus modulating hippocampal function by altering chromatin accessibility and transcriptional pathways.

Since free serotonin levels in the murine hippocampus are ∼300 fM (Schechter et al., 2007) while the K_D_ for HTR6 binding to serotonin is 12.6 nM (Boess et al., 1997), the axo-ciliary synapses provide a mechanism to localize and concentrate serotonin’s effect. The ciliary HTR6-Trio-RhoA signaling axis limits serotonin-RhoA signaling to the cilium and exploits its specialized link to the nucleus, much as the Shh pathway regulating Gli transcription factors is limited by compartmentalized G_αs_/PKA signaling. This rationalizes the persistence of primary cilia in non-dividing mature cells such as neurons and can explain how alterations in ciliary signaling can impact structures such as excitatory synapses on dendrites that can be hundreds of microns distant (Tereshko et al., 2021). Interestingly, in other cells such as pre-adipocytes, omega-3 fatty acids were shown to activate primary ciliary FFAR4 receptors to induce CTCF-dependent chromatin changes through opening of CTCF-binding enhancer sites (Hilgendorf, 2019). This posits a tantalizing theme in which primary cilia act as an epigenetic regulator to stabilize transcriptional programming in response to environmental cues. In this “cilia as the nuclear antenna” model, cilia provide a shorter and more direct spatial pathway for neurotransmitters and other receptor agonists to specifically regulate nuclear transcription.

Not all cilia form serotonergic axo-ciliary synapses. This raises the question of whether the neuron hosts of these cilia are specialized among CA1 pyramidal neurons. Perhaps the non-serotonergic synapsing cilia are enriched in other GPCRs (Dopamine receptor 1 (DRD1), somatostatin 3 (SSTR3), galanin receptors (Hilgendorf et al., 2016) or tyrosine kinase receptors that forms axo-ciliary synapses with a different set of axons and elicit different functional networks. The fact that the portion of cilia with axo-ciliary synapses detected in FIB-SEM (∼67% to 80%) is greater than those found with HTR6-serotonergic axons (35%) supports this possibility. Further studies examining different ciliary receptor and axon types will address these questions.

### Ciliary signaling and chromatin remodeling

It is not currently clear how the loss of serotonin mediated ciliary RhoA activation results in changes in nuclear actin and chromatin accessibility. Presumably, activated RhoA may directly traffic to the nucleus, but ciliary RhoA activation may activate Rho-associated kinases (ROCK1 and ROCK2), which subsequently localize to the nucleus. Nuclear ROCK2 has been shown to phosphorylate EP300, or histone acetyltransferase p300, that can regulate transcription via chromatin opening (Tanaka et al., 2006). Our current data suggests that serotonin signaling in adult primary cilia maintains one state, and its removal initiates a change leading to adducin/actin mediated chromatin remodeling.

The time variation of physiological changes in brain serotonin provokes other questions. The raphe serotonergic system is much more active in wake states than during sleep (Oikonomou et al., 2019; Wan et al., 2021). Interestingly, the transcript levels of HTR6 also oscillates and peaks around midnight, or before the serotonergic system becomes active (Baldi et al., 2021). We do not know if HTR6 axo-ciliary synapses oscillate and may be involved in chromatin remodeling during sleep/wake cycles (Hor et al., 2019) which may impact learning and memory (Rasch and Born, 2013). A recent genome-wide association study identified HTR6 as one of the 15 genes indicated in bipolar disorders (Mullins et al., 2021), while a somatic HTR6 mutation was detected in resected cortical seizure foci with dysplastic growth (Zhang et al., 2020). In addition, HTR6 is also expressed in upper and lower motor neurons, another class of pyramidal neurons (Bandyopadhyay et al., 2013). Interestingly, in zebrafish, serotonergic inputs to the motor neurons promote adult regeneration of motor neurons (Barreiro-Iglesias et al., 2015). It will be interesting to determine if serotonergic axo-ciliary synapse and HTR6-RhoA driven chromatin maintenance are also present in these neurons, which may provide mechanistic insight into their survival.

The major importance of this work is the discovery of a new kind of chemical synapse, one that conveys information more directly to the target cell’s nucleus. A remaining question is whether the axon-cilia-nucleus axis is a general mechanism for maintaining a stable state in neurons and other cells, as loss of this function results in major chromatin reorganization. The current work suggests that loss of ciliary HTR6 RhoA activation results in nuclear actin changes and alterations in chromatin accessibility. Finally, does disruption of these primary cilia-dependent stable connections lead to programmatic functional changes, resulting in changes in memory or, for example, differentiation status? More broadly does disruption of primary cilia-specific receptors contribute to cancer, type II diabetes, and other disease states in brain and other organs?

## Acknowledgements

We thank members of the Clapham Lab, Lavis Lab, Lippincott-Schwartz Lab, and the Marin Lab for productive discussions and critical reading of the manuscript (Drs. William Valinsky and Alex Miller). We thank Liangqi (Frank) Xie (Tjian and Liu Lab) for kindly providing the Tn5 transposase and Tn5 transpose-ATTO-590-oligonucleotide mixture solution. We thank Dr. Joachim Goedhart (University of Amsterdam) and Dr. Kees Jalink (Netherlands Cancer Institute) for input on our RhoA-FRET and RhoA-FLIM analyses, respectively. We thank Jeffrey Marshman (Zeiss) for help with the Zeiss Crossbeam FIB-SEM. We thank the Janelia Anatomy and Histology, Cell and Tissue Culture, Vivarium, Molecular Biology Shared Resources, and iEplore-Biocampus animal facility for their support. **Funding:** This work was supported by the Howard Hughes Medical Institute. The generation of Htr6-EGFP knock-in mice and Htr6 KO mice were funded by the Foundation for Medical Research (FRM, France), and two ANR contracts Sero6Cognet (ANR-11-BSV4-008), and Sero6Dev (ANR -17-CE16-0010-01). V.D. was supported by the French ministry of research and education. S.H.S. was supported by NIH 5T32HL110852-05. S.E.B. is supported by ANID-Millennium Science Initiative Program #NC160011. S.U. was supported by Philomathia Foundation and Chan Zuckerberg Initiative Imaging Scientist program. T.K. was supported by National Institute of General Medical Sciences Maximizing Investigators’ Research Award GM130386; National Institutes of Health R01 GM075252; Biogen Sponsored Research Agreement.

## Author Contributions

Conceptualization, S.H.S. and D.E.C.; Methodology, S.H.S., S.E.B., S.B., C.S.X, F.D., J. W., Y.L.L, J.H., T.K., A.L.L.; Investigation, S.H.S., G.U., V.D., S.E.B., S.B., A.L.L., L.W., F.D., J.W., Y.L.L, M.F., H.A.P., S.P., C.S.X., T.K.; Writing – Original Draft, S.H.S.; Writing -Review & Editing, S.H.S., L.L., S.E.B., S.C.D., D.E.C.; Funding Acquisition, S.H.S., L.L., S.C.D., T.K., and D.E.C.; Resources, H.F.H.; Visualization, S.H.S., G.U., A.L.L.; Supervision, S.H.S. and D.E.C.

## Supplementary Figure Legends

**Table S1. Number of WT and KO cells in each individual cluster and marker genes.**

Related to Figure 7.

**Video S1**. **Reconstruction of CA1 neuronal cilia.** Related to Figure 2. Movie showing the reconstructed CA1 pyramidal neuronal primary cilia using 8-nm isotropic FIB-SEM imaging.

## STAR METHODS

### RESOURCE AVAILABILITY

#### Lead contact

Further information and requests for resources and reagents should be directed to and will be fulfilled by the lead contact, David Clapham.

#### Materials availability

All unique/stable reagents generated in this study are available from the lead contact. Sharing of the RPE-1 cells are limited by terms set by ATCC.

#### Data and code availability

scATAC-seq datasets have been deposited to Gene Expression Omnibus (accession number GSE183431).

Any additional information required to reanalyze the data reported in this paper is available from the lead contact upon request.

### EXPERIMENTAL MODEL AND SUBJECT DETAILS

#### Animals

All animal work was approved by the Boston Children’s Hospital Institutional Animal Care and Use Committee (IACUC 16-03-3138R), the Janelia Institutional Animal Care and Use Committee (IACUC 16–146 and 19-181), or the animal use and care guidelines of Montpellier University (France, authorization D34-172-4). C57BL/6 mice were obtained from Charles River Laboratories. Htr6-EGFP knock-in mice were generated at the Institut Clinique de la Souris (Illkirch-Graffenstaden, France). Htr6 KO mice were generated at the Phenomin consortium (Institut Clinique de la Souris, Illkirch-Graffenstaden, France) by using CRISPR-Cas9. Htr6 exon 3 and 4 were targeted using two pairs of guide RNAs on each side of the targeted region. No restrictions were imposed on food and water. For doxycycline induction experiments, mice were fed with doxycycline-containing food (2000 ppm, Animal Specialties and Provisions, modified from LabDiet 5053) for 1 week. Similar results were obtained in both males and females.

#### Cell culture

RPE-1 (female) and HEK293A cells (gift from Dr. Asuka Inoue, Tohoku University, Japan) were plated at ∼20,000 cells/cm^2^ on #1.5 12 mm coverslips in 24-wells, 1-chamber 35 mm glass-bottom dishes, or 4-chamber 35 mm glass-bottom dishes (all #1.5 cover glass, Cellvis) in 10% serum containing media (RPE-1: DMEM:F12 media, ATCC 30-2006; HEK293A: DMEM, low glucose, GlutaMax, pyruvate, Thermo Fisher Scientific #10567014; Day 0). With HEK293A cells, the dishes were coated with Matrigel (Corning Life Sciences) before plating. The next day (Day 1), the cells were serum deprived with 0% FBS media with 100 ng/ml doxycycline to induce GRAB-HTR6-cilia, HTR6-RhoA, or Arl13b-RhoA expression. For HEK393A cells, 1 μM H-89 was also added to induce ciliogenesis. After 24 h of serum deprivation (Day 2), GRAB-HTR6-cilia cells were labeled with 250 nM Janelia Fluor 552 (JF552) dye for 2 h. Cells were rinsed and placed back in serum-free media without doxycycline or H-89. Experiments were conducted at 48 to 96 h after serum deprivation (Day 3 to 5).

For stable cell line creation, hTERT RPE-1 cells (ATCC CRL-4000) or HEK293A cells were transfected with the piggyBac hyperactive transposase vector (VectorBuilder) and HTR6-RhoA sensor vector or GRAB-HTR6-cilia concurrently with Lipofectomine 3000 (Thermo Fisher Scientific) at a 1:2.5 ratio and grown in 10% Tet-free FBS (Gemini) containing media. The cells were then selected by blasticidin at 10 μg/ml to create Tet-on HT6-RhoA sensor stable cells.

#### Primary hippocampal and raphe neuron culture

Hippocampi and midbrains were dissected from P0 Sprague-Dawley rat pups of both sexes. Rat maintenance and care followed policies advocated by NRC and PHS publications and approved by the Institutional Animal Care and Use Committee (IACUC), Janelia Research Campus. Tissues were digested with papain and gently triturated and filtered through a 40 µm filter. Neurons were electroporated (Lonza 4D-nucleofactor) with Arl13b-RhoA (hippocampal neurons) or Tph2-Cre (tryptophan hydroxylase 2-Cre, midbrain neurons) and plated in poly-D-lysine coated dishes and cultured in NbActive medium (Brainbits) at 37°C and 5% CO_2_. A week after plating, the neuronal cultures were fed with B-27 plus neuronal culture system (Thermo Fisher Scientific). AAV transduction (FLEX-on hM3DGq-DREADD, FLEX-on farnesylated SNAP, or FLEX-on tdTomato) were applied at DIV10. 100 ng/ml doxycycline was added at DIV 6 or DIV14 and removed the next day. Images were collected between DIV 9 and DIV 28 (Alr13b-RhoA, hippocampal culture only) or DIV21-35 (hippocampal and midbrain co-culture). JF552-STL (SNAP-labeled neurons) was applied the day prior to imaging.

### METHOD DETAILS

#### Synthesis of PA-O-5-hydroxytryptamine (photoactivatable serotonin)

*N*-Boc serotonin (**S1**, 60 mg, 217 µmol, 3.5 eq) and coumarin bromide (**S2**, 30 mg, 62.2 µmol, 1 eq) were dissolved in CH_3_CN (4 mL). K_2_CO_3_ (potassium carbonate, 60 mg, 435 µmol, 2 eq) was added and the reaction was stirred at room temperature for 15 h. The reaction was concentrated under reduced pressure and the residue was dissolved in EtOAc. This was washed with water and saturated NaCl (aq), dried over MgSO_4_, and concentrated under reduced pressure. The material was purified using flash chromatography on silica gel (0–50% EtOAc/hexanes, linear gradient), which afforded 35 mg (83%) of compound **S3** as a pale-yellow solid. Compound **S3** (30 mg, 44.3 µmol) was dissolved in CH_2_Cl_2_ (2 mL). Trifluoroacetic acid (TFA; 0.4 mL) was added and the reaction was stirred at room temperature for 2 h while shielded from light. Toluene (5 mL) was added and the mixture was concentrated under reduced pressure. The residue was purified by reverse-phase HPLC using a gradient of CH_3_CN/H_2_O containing 0.1% v/v TFA as additive. 1H NMR (400 MHz, 1:1 CD3OD, CD3CN) δ 7.71 (s, 1H), 7.64 (d, J = 8.9 Hz, 1H), 7.31 (d, J = 8.9 Hz, 1H), 7.16 (d, J = 2.4 Hz, 1H), 7.13 (s, 1H), 6.93 (dd, J = 8.8, 2.5 Hz, 1H), 6.65 (dd, J = 9.0, 2.7 Hz, 1H), 6.52 (d, J = 2.6 Hz, 1H), 6.32 (s, 1H), 5.30 (s, 2H), 4.24 (s, 4H), 3.15 (t, J = 7.3 Hz, 2H), 3.01 (t, J = 7.3 Hz, 2H). HRMS (ESI) calculated for C_24_H_24_N_3_O_7_ [M+H]+ 466.1609, was 466.1615.

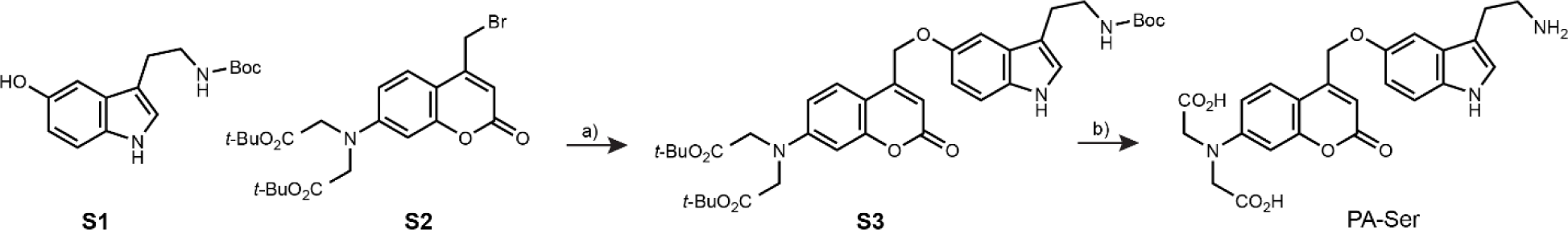

#### Molecular Biology

The piggyBac Tet-On HTR6-RhoA sensor was generated based on the published mScarlet-I based RhoA sensor (Bindels et al, 2016). The mScarlet-I:sGFP2:RhoA and the cpPKN1 fragments were synthesized (Genscript) and cloned into piggyBac Tet-On vector Xlone HTR6-HaloTag (Deo et al., 2019). The Xlone-Ar13b-RhoA sensor was subcloned by replacing HTR6 with ARL13B (VectorBuilder). GRAB-HTR6-cilia was made by removing the IgG leader sequence and adding a HaloTag to the C-terminus based on GRAB-HTR6-PM (see Figure S3 and below) by VectorBuilder. Tph2-Cre vector was generated by replacing GFP in Tph2-GFP (Wan et al, 2016; gift from Dr. Björklund) with the Cre-recombinase in Syn1-EBFP-Cre AAV (gift from Hongkui Zeng, Addgene plasmid # 51507; RRID:Addgene_51507). pAAV-Syn-FLEX-rc[ChrimsonR-tdTomato] was a gift from Edward Boyden (Addgene plasmid # 62723; RRID:Addgene_62723). pAAV[FLEX-ON]-CAG-Farnesylated-SNAPtag AAV plasmid was made by VectorBuilder, Inc. pAAV-FLEX-ON-tdTomato was a gift from Edward Boyden (Addgene plasmid # 28306; RRID:Addgene_28306). pAAV-EF1a-DIO-hM3D(Gq)-mCherry was a gift from Bryan Roth (Addgene plasmid # 50460; RRID:Addgene_50460). pAAV-TRE-HTR6-SNAPf- TRIP and pAAV-Syn-Tet3G were made by VectorBuilder. Syn-Tet3G and TRE-HTR6-SNAPf-TRIP were packaged in AAV2/rh10 capsid, and the other viral vectors were packaged in the AAV2/PHP.eB capsid. All viral vectors were made by the HHMI Viral Tools.

#### Intracranial injections

6-week-old adult mice were anesthetized with 2.5%-3.0% isoflurane with an O_2_ rate of 1L/min and mounted on a stereotaxic frame. The body temperature was maintained at 37 °C using a heating pad. Mice were given buprenorphine (0.1mg/kg) subcutaneously at 5mg/kg of body weight. After shaving, a drill was used to create a small craniotomy hole (∼1 mm). 200 nL of AAV mixture composed of AAV-rh10-Syn-Tet3G and AAV-rh10-TRE-HTR6-TRIP (1:2 ratio, 5×10^12^ vg/ml and 10^13^ vg/ml, respectively) were bilaterally injected in the mouse CA1 pyramidal neuron layer (anteroposterior –2.25 mm relative to Bregma, mediolateral ±2 mm relative to Bregma; dorsoventral –1.43 mm relative to the pial surface) at a rate of 50 nl/minute using a Nanoject II Injector (Drummond Scientific, USA) followed by 5 additional minutes to allow diffusion. Upon recovery, mice were given Ketaprofen (5mg/kg, sub-cutaneous) and placed back in the home cage. Ketoprofen (5mg/kg, sub-cutaneous) was given once a day for an additional two days post-surgery.

#### Immunofluorescence and Imaging

For cultured cells, #1.5 coverslips were first fixed in 4% paraformaldehyde (PFA) in PBS overnight at 4°C. Samples were then rinsed in PBS for 5 min x 3. After permeabilization with 0.3% Triton-X in PBS for 30 min, samples were blocked with 5% normal goat serum (NGS) in PBS for 1 h. 5% NGS was then replaced with primary antibody containing solution in 1% BSA in PBS with 2 mM sodium azide overnight at 4°C. After rinsing in PBS for 5 min x 3, samples were stained with secondary antibody and Hoescht 33342 for 2 h. For Trio staining, the samples were stained according to the Alexa tyramide amplification system with the SuperBoost protocol (Thermo Fisher Scientific). Samples were then rinsed in PBS for 5 min x 2 /Milli-Q water 5 min x 1 and mounted on Vectashield Vibrance (Vector Laboratories) or Prolong Fade Glass (Thermo Fisher Scientific).

For mouse brain samples, mice were deeply anesthetized with isoflurane inhalation or ketamine/xylazine (200 mg/kg ketamine and 20 mg/kg xylazine) intra-peritoneal injection and intracardially perfused with 4% PFA in phosphate-buffered saline (PBS). The brains were removed and post-fixed in 4% PFA in PBS overnight at 4°C. Samples were then rinsed in PBS x3, 15 min each. Serial sections of 50 μm or 200 μm were obtained on a vibratome. For 50 μm mouse brain sections, fixed slices were incubated at room temperature in 0.3% Triton-X in PBS overnight, followed by 5 h in blocking buffer (BlockAid, Thermo Fisher Scientific), overnight in primary antibody, and again overnight in secondary antibody and Hoechst 33342 (2 μg/ml). Both primary and secondary antibodies were diluted in the same blocking buffer (BlockAid, Thermo Fisher Scientific). The stained sections were then rinsed in PBS 15 x 2 times/300 mOsm glycerol 15 min x 1, and mounted in Prolong Fade Glass (Thermo Fisher Scientific). Between antibodies, slices were washed x3, 15 min each with PBS. For 200 μm mouse brain sections, slices were first treated with the epoxide-crosslinker as in the SHIELD protocol (Life Canvas Technology, Park et al, 2018). Afterwards, sections were incubated in 2% Triton-X in PBS with 2 mM sodium azide for 3 days for permeabilization. Sections were then incubated in primary and secondary antibodies for 2 days at room temperature, respectively. After washing, sections were mounted in SlowFade Glass (Thermo Fisher Scientific).

The following primary antibodies were used: rabbit anti-GFP (Chromotek PABG1, 1:1000), guinea-pig anti-SERT (synaptic systems 340004, 1:200), mouse anti-ADCY3 (Encor Biotechnology MCA-1A12, 1:1000), rabbit anti-PCP4 (Millipore Sigma, HPA005792, 1:500), chicken anti-rootletin (Millipore Sigma, ABN1686, 1:1000), rabbit anti-ADD1 (Abcam EP734Y, #ab40760, 1:250), rabbit anti-synaptophysin (Cell Signaling, #36406, 1:100), rabbit anti-synaptophysin (Thermo Fisher Scientific, MA5-14532, 1:200), rabbit anti-Trio GEFD2 (custom antibody, gift from Dr. Susanne Schmidt, 1:500). The following secondary antibodies and dyes were used: Alexa Fluor Plus 488 goat anti-rabbit (1:1000, Thermo Fisher Scientific), Alexa Fluor 488 donkey anti-rabbit (1:400, highly-cross absorbed, Jackson ImmunoResearch), CF488 donkey anti-guinea pig (1:1000, Biotium), Alexa Fluor Plus 555 goat anti-rabbit (1:1000, Thermo Fisher Scientific), CF633 goat anti-mouse IgG_1_ (1:1000, Biotium), Hoescht 33342 (2 μg/ml, Thermo Fisher Scientific), SuperBoost goat anti-rabbit polyHRP (ready-to-use 1x concentration, Thermo Fisher Scientific).

All confocal imaging employed a Zeiss 880 Laser Scanning Confocal Microscope (LSM) equipped with a Plan-Apochromat 63x oil objective (Zeiss, NA = 1.40), 40x multi-immersion LD LCI Plan-Apochromat objective (Zeiss, NA = 1.20), or 20x air Plan-Apochromat objective (Zeiss, NA = 0.8); ZEN Black software (Zeiss). Hoechst 3342 was excited by 405 nm laser light and the spectral detector set to 409-481 nm. Alexa 488/CF488 was excited by 488 nm laser light and the spectral detector set to 490-545 nm. Alexa 555 was excited by 561 nm laser light and the spectral detector set to 570-642 nm. ATTO-590 was excited by 594 nm laser light (Airyscan only). CF633 was excited with 633 nm light and the spectral detector set to 642-755 nm. The spectral detector was only used for non-Airyscan confocal scanning imaging sessions.

### Sample preparation, FIB-SEM imaging, and data analyses of mouse brain samples

#### Conventional chemical fixation protocol

Mice were anesthetized with ketamine/xylazine (200 mg/kg ketamine and 20 mg/kg xylazine) intra-peritoneal injection and perfused with a solution of 2% PFA and 2% glutaraldehyde in 0.1 M sodium cacodylate buffer, 0.2 mM CaCl_2_. The brain samples were dissected and post-fixed in the perfusion solution overnight at 4°C. After rinsing with 0.1M sodium cacodylate buffer, 300 μm serial sections were obtained on a vibratome. The tissue was then immersed in 1% osmium tetroxide and 1.5% potassium ferricyanide in a 0.1 M cacodylate buffer for 1 h. After rinsing in 0.1 M sodium cacodylate buffer, the sections were further stained with 1% osmium tetroxide in water for 1 h, followed by 2% uranyl acetate in maleate buffer (pH = 5.15) overnight at 4°C. The tissue was then washed in water, dehydrated with graded ethanol, and embedded in Epon812 resin.

Epon-812 flat-embedded mouse hippocampal CA1 samples were first mounted on an aluminum stub. The sample surface was polished on an ultramicrotome, followed by carbon coating (20 nm). The samples were then imaged on the Zeiss Crossbeam 540 at 5-6 nm pixel size with 15-20 nm milling using ATLAS 5 software (Zeiss).

#### Hybrid protocol

2- to 3-month-old male C57/BL6 mice were deeply anesthetized and transcardially perfused with 30 mL of 3% PFA (60 mM NaCl, 130 mM glycerol, 10 mM sodium phosphate buffer). The brain was carefully dissected from the skull and post-fixed with 50 mL of 3% PFA (30 mM NaCl, 70 mM glycerol, 30 mM PIPES buffer, 10 mM betaine, 2 mM CaCl_2_, 2 mM MgSO_4_) at room temperature for 2 h. The brain sample was then rinsed in a 400 mOsM buffer (65 mM NaCl, 100 mM glycerol, 30 mM PIPES buffer, 10 mM betaine, 2 mM CaCl_2_, and 2 mM MgSO_4_) for 0.5 h, followed by vibratome sectioning (coronal sections, 100 μm thickness) using a Leica VT1000S vibratome in the same buffer. 100 μm sections were then fixed in 1% PFA, 2% glutaraldehyde solution (30 mM NaCl, 70 mM glycerol, 30 mM PIPES buffer, 10 mM betaine, 2 mM CaCl_2_, 2 mM MgSO_4_, 75 mM sucrose) overnight at 4°C. Sections were then washed using the 400 mOsM rinsing buffer (see above). Round samples of the hippocampus were created from the 100 μm coronal sections using a 2 mm biopsy punch (Miltex). The 2 mm samples were dipped in 1-Hexadecene, placed in a 100 μm aluminum carrier, covered with a flat carrier and high-pressure frozen using a Wohlwend compact high-pressure freezer (Wohlwend GmbH, Switzerland). Samples were then freeze-substituted in 0.5% osmium tetroxide, 20 mM 3-amino-1,2,4-triazole or 20 mM imidazole, 0.1% uranyl acetate, 4% water in acetone, using a Leica AFS2 system. Specimens were further dehydrated in 100% acetone and embedded in Durcupan resin.

Two datasets were acquired using the hybrid protocol, stained with either osmium-imidazole or osmium-3-amino-1,2,4-triazole. The samples were mounted on a copper post and trimmed to the Region of Interest (ROI), guided by X-ray tomography data obtained by a Zeiss Versa XRM-510. The samples were coated with a thin layer of 10- to 20-nm gold and 50- to 100-nm carbon and imaged by a customized Zeiss Merlin FIB-SEM or NVision40 FIB-SEM system using 8nm pixel size with 2 or 4 nm of milling depth. After alignment using a Scale Invariant Feature Transform (SIFT) based algorithm (Lowe, 2004), the stacks were binned by a factor of 2 or 4 along z to form a final isotropic volume of 35 µm x 35 µm x 40 µm and 50 µm x 50 µm x 44 µm with 8 nm x 8 nm x 8 nm voxels.

The electron microscopy datasets generated above were manually segmented using VAST (Volume Annotation and Segmentation Tool, Berger et al., 2018). Segmented results were exported as obj files, and rendered using 3ds Max 2021 (Autodesk, Inc.). For synaptic vesicles in Figure 2E, the centroids of segmented vesicles were calculated in 3D, and 40 nm spheres were generated as 40 nm spheres in 3ds Max.

The hybrid protocol was developed for this work, which were used to generate two enhanced FIB-SEM datasets in the CA1 area to examine primary cilia. Prior to the completion of this manuscript, these datasets have been shared with others to examine lipid droplets (Ioannou et al., 2019), myeline distribution along large axons (Gao et al., 2019), and mitochondrial morphology (Thomas et al., 2019). All FIB-SEM datasets will be available upon request.

### Characterization of GRAB-HTR6-PM sensor

#### Expression of the GRAB-HTR6-PM sensor in HEK293T cells

HEK293T cells were cultured in DMEM (Gibco) supplemented with 10% (v/v) FBS (Gibco) and 1% penicillin-streptomycin (Gibco) at 37℃ in 5% CO_2_. HEK293T cells were plated on 96-well plates and transfected with a mixture of plasmids and PEI (300 ng plasmids and 900 ng PEI for each well) when the cells were grown to ∼70% confluence. The medium was replaced after 4-6 h, and cells were used for imaging 24 h after transfection.

#### Fluorescence imaging of HEK293T cells expressing the GRAB-HTR6-PM sensor

The DNA for the IRES-mCherry-CAAX cascade was fused downstream of the GRAB-HTR6-PM sensor to normalize the expression level of the sensor and calibrate the membrane signal. HEK293T cells expressing the GRAB-HTR6-PM sensor were imaged by the Opera Phenix high-content screening system. Before imaging, the culture medium was replaced with 100 μl Tyrode’s solution. For imaging, a 40x, 1.1-NA water-immersion objective, a 488-nm laser combined with a 525/50-nm emission filter for excitation and collection of the GFP fluorescence signal, and a 561-nm laser combined with a 600/30-nm emission filter for the collection of the mCherry fluorescence signal were used. The same field of views (FOVs) were imaged without or with the application of 5-HT (at various concentrations, in Tyrode’s solution), respectively. The fluorescence signal of the GRAB-HTR6-PM sensor was calibrated using the ratio of GFP to mCherry.

#### Quantification and statistical analysis of GRAB-HTR6-PM sensor

Images collected by an Opera Phenix high-content screening system from cultured HEK293T cells were processed using ImageJ (1.53c) software (NIH) and analyzed using custom-written MATLAB (R2020b) codes. The fluorescence response (ΔF/F_0_) was calculated using the formula (F−F_0_)/F_0_, in which F_0_ is the baseline fluorescence signal after subtracting the background. The dose-response curve was plotted using OriginPro (2020b).

### Measurements of the GRAB-HTR6-cilia sensor

#### GRAB-HTR6-cilia titration curve

Serum deprived, JF552 labeled RPE-1 cells stably expressing GRAB-HTR6-cilia were imaged in a HEPES-buffered imaging media (140 mM NaCl, 20 mM HEPES, 2.5 mM KCl,

1.8 mM CaCl_2_, 1.0 mM MgCl_2_, pH = 7.4, mOsm = 300; Live Cell Imaging Solution, Thermo Fisher Scientific A14291DJ) using a 20x air Plan-Apochromat objective (Zeiss, NA = 0.8) with FAST Airyscan on a Zeiss 880 confocal microscope at 37°C. GFP and JF552 were excited with 488 nm and 561 nm lasers, respectively. Keeping the same field of view, Z stacks were acquired at different concentrations of 5-HT diluted in the same imaging buffer. The two channels were aligned using Zen Blue (Zeiss). Cilia were segmented using CiliaQ based on the JF552 channel (Hansen et al., 2021). Per cilium GFP mean fluorescent intensities were then calculated for different concentrations. The fluorescence response (ΔF/F_0_) was calculated using the formula (F−F_0_)/F_0_, in which F_0_ is the baseline fluorescence signal after subtracting the background. The dose-response curve was plotted using Graphpad Prism 9.2.

#### GRAB-HTR6-cilia in hippocampal-midbrain neuron co-culture

Hippocampal neurons expressing GRAB-HTR6-cilia and serotonergic neurons expressing ChrimsonR-tdTomato, farnesylated SNAP-Tag:JF552, or tdTomato were imaged with a 40x multi-immersion LD LCI Plan-Apochromat objective (Zeiss, NA = 1.2; silicone oil was used as the immersion media) on a Zeiss LSM 880 microscope in artificial cerebral spinal fluid (NaCl 124 mM, KCl 2.5 mM, NaH_2_PO_4_ 1.2 mM, NaHCO_3_ 24 mM, HEPES 5 mM, glucose 12.5 mM, MgSO_4_ 2mM, CaCl_2_ 2mM, Ting et al., 2018) at 37°C. Two-channel FAST Airyscan images were first imaged to characterize the axociliary synapses using 488 nm and 561 excitations for GFP and red fluorophores, respectively. GRAB-HTR6-cilia was then imaged at 1 Hz using the 488 nm laser with Z-stacks. After 30 s (frame = 31), the 594 nm laser line was used to photostimulate at the same 1 Hz frequency (repetition = 1, same pixel dwell time as imaging) as image acquisition (488 nm). A total of 120 Z-stacks were acquired per cilium (120 s). Airyscan stacks were processed using Zen Black (auto-strength, 3D; Zeiss). These 4D stacks (XYZT) were projected onto single planes (XYT) by maximum intensity projection in Fiji/ImageJ. Cilia were segmented using CiliaQ (Hansen et al, 2021). The first 10 time points, which often showed significant quenching, were discarded for subsequent analyses. A 1-μm circle at the contact site (ChrimsonR, tdTomato, or SNAP:JF522) or at the site at which the cilium is closest to a serotonergic axon was used as the region of interest (ROI) to calculate mean intensity values over time (ImageJ/Fiji). These values were processed with a low pass filter at 0.2 Hz. The average intensity of the 10 time points prior to photostimulation was used as the baseline fluorescence intensity, or F_0_. The maximum ΔF/F_0_ during photostimulation was used for statistical comparisons in different conditions using estimation statistics (Ho et al., 2019). Low pass filter, baseline fluorescence intensity calculation, and estimation statistics were done in a custom Python code.

### HTR6-RhoA FRET measurements and analysis

Images were collected using silicone-oil immersion media with a 40x multi-immersion LD LCI Plan-Apochromat objective (Zeiss, NA = 1.2; silicone oil immersion media) on a Zeiss LSM 880 microscope. Single apical cilia were imaged at zoom 10 with 4 Airy units (146 µm pinhole), 212 x 212 frame size, 0.1 µm pixel size, 0.93 µs pixel dwell time, 16-bit bidirectional scanning, and 4 optical sections (1.8 µm) to cover the entire length of the cilium. The resulting temporal resolution was 0.25 s. The donor fluorophore, sGFP2 was excited by the 488 nm laser. Donor emission from mScarlet-I was collected with the spectral detector set to 490-550 nm. Acceptor-sensitized emission was collected with the spectral detector set to 570-650 nm.

For uncaging experiments, the following uncaging parameters were used on a 5 µm radius circle adjacent to the ciliary tip: repeat each stack x10 using the same pixel dwell time as during scanning. Two experiment blocks (400 timeframes each) were used to acquire data with and without uncaging. Uncaging started at frame 20 in the first block, with no uncaging during the second block.

The 4D stacks (XYZT) were first projected onto single planes (XYT) by maximum intensity projection in Fiji/ImageJ. Datasets with bidirectional scanning artefacts (i.e. misalignment between two scanning directions in alternating lines), significant movements, and/or focus drifts were excluded. The donor channel was used to create a mask to segment the cilia (“cilia mask”) by using the “subtract background with smoothing paraboloid” command followed by Otsu thresholding in Fiji/ImageJ. The donor and sensitized emission intensity values were individually subtracted by using the mean intensity from an acellular area outside the cilium. These background-subtracted images were then masked using the cilia mask to isolate/segment cilia-specific signals. The FRET ratio was calculated by dividing the segmented sensitized emission by segmented donor emission. For direct ligand application experiments, FRET recordings were digitized at 4 Hz and filtered at 1 Hz with a low pass Fourier transform digital filter implemented in Origin (OriginLab Corporation). For uncaging experiments, the FRET data was processed with temporal averaging by a factor of 2. Change of the mean FRET ratio of the entire cilium was plotted using the Z plot function in Fiji/ImageJ and imported into Excel spreadsheets. The changes were visualized using PlotTwist (Goedhart, 2020). ΔF/F was calculated by using the mean FRET ratio at the beginning of the experiments as the baseline (frame 1 to 25 for direct ligand application, and 1 to 30 for uncaging experiments). Statistical testing of RhoA spikes were done in Prism 8 (GraphPad).

### Ciliary-RhoA FLIM measurements and analysis

3D Z stacks were collected using an 40x oil immersion Plan-Apochromat objective (Leica, NA = 1.3) on a Leica SP8 Falcon microscope at 2x the Nyquist limit and 1 Airy disc. Donor sGFP fluorescence was excited by a pulsed (40 MHz) white light laser tuned at 488 nm, and emitted photons between 490 nm and 550 nm were collected with line repetition = 8. To calculate the fluorescence lifetime of unquenched and quenched donor, large populations of cells expressing the RhoA sensor were imaged, and fluorescence lifetimes were calculated by the n-exponential reconvolution fitting algorithm (2 exponential components) with pixel binning by a factor of 2 (Leica LAS X FLIM/FCS software v3.5). The fluorescence lifetime of quenched and unquenched donor was determined to be 1.3 and 2.7 ns, respectively. These numbers were then used for all subsequent fittings. Each cilium is imaged individually with Z-stacks. To calculate representative RhoA sensor sGFP fluorescence lifetime per cilium, “FLIM Image Fit” was performed in the Leica LAS X FLIM/FCS software suite. The resulting datasets were rendered in 3D for visualization (Figure 5A, B, D, F, G, I, and J) and exported as two-channel stacks, encoding photon counts and fluorescence lifetime, respectively. Subsequent imaging analyses were done in Python. For each stack, the channel encoding counts were used to segment cilia (Otsu and Yen thresholding were used for HEK cells and neurons, respectively). The segmented cilia were then used as masks to extract ciliary voxels from the FLIM channel. Voxels with less than 50 counts were discarded. An alpha distribution was fitted to the histogram of the FLIM channel, as it provided the best fitting among all 80 different statistical distributions tested. The peak of the alpha distribution was used as the mode, or the representative FLIM of a given cilium. Statistical comparisons of cilia from different cell types and states, and after stimulation were performed using estimation statistics (Ho et al., 2019).

### 3D ATAC-Airy staining and analysis

Transposase-mixture solution (hyperactive Tn5 transposase with ATTO-590 or ATT-565 conjugated oligos) were prepared as described previously (Chen et al., 2016). 50 μm thick, PFA-fixed CA1 coronal sections from 6-month-old C57BL/6 wild-type, HTR6 KO, non-doxycycline-treated control and doxycycline-treated HTR6-TRIP mice were obtained using the same method as described above. A 3 mm punch of the CA1 area was created using a biopsy punch on the 50 μm thick sections (Electron Microscopy Sciences). These punches were permeabilized in 0.3% Triton/PBS overnight at RT and blocked with buffer (BlockAid, Thermo Fisher Scientific) for 1 h at RT. The samples were then rinsed in PBS for 15 min x 3 and incubated in the transposase-mixture solution (100 nM, 10 mM Tris-HCL pH 7.5, 10 mM MgCl_2_, 25% dimethylformamide) overnight at 4°C, followed by 1 h at 37°C on a rotator. After incubation, the samples were washed x 3 for 15 min at 55 °C with 1×PBS containing 0.01% SDS and 50 mM EDTA, followed by regular PBS at room temperature for 15 min. The samples were subsequently stained with Hoechst 33342 (2 ug/ml) for 2 h, rinsed in PBS 15 x 2/300 mOsm glycerol x1, and mounted in Prolong Fade Glass (Thermo Fisher Scientific). Non-doxycycline-treated control and doxycycline-treated HTR6-TRIP samples were also stained with ADCY3 and CF633 (see above).

For quantification of ATAC-see intensity per nucleus, CA1 areas were imaged using a 40x multi-immersion LD LCI Plan-Apochromat objective (Zeiss, NA = 1.2; doxycycline-treated control and doxycycline-treated HTR6-TRIP) or a 63x Plan-Apochromat oil-immersion objective (WT and KO) using FAST Airyscan or Airyscan, respectively. The FAST Airyscan and Airyscan stacks were processed with 3.4 and 6.7 filter strength/3D mode in Zen Black or Zen Blue, respectively (Zeiss; voxel size 41 nm x 41 nm x 1939 nm and 35 nm x 35 nm x 144 nm). The stacks were then downsampled to a voxel size of 400 nm x 400 nm x 400 nm and imported into Cellpose (Stringer et al., 2021). The Hoechst channel was used to segment each individual nucleus using “cyto2”. Segmentations less than 5000 voxels (equivalent to a sphere with an 8-µm diameter) were excluded, and the mean intensities of the nuclei were calculated in a custom Python code for statistical analyses by estimation statistics (Ho et al, 2019).

### Single Cell ATAC-seq

One frozen hippocampus each from KO and wild-type mice were prepared for scATAC-seq according to the manufacturer’s protocol for flash frozen mouse brain (Protocol 2.2, CG000212 Rev B, 10X Genomics). Nuclei were counted using a Luna-II automated cell counter (Logos Biosystems). Approximately 12,000 nuclei per sample were subjected to the Chromium NextGem scATAC-seq v1.1 assay (10X Genomics) and the resulting libraries were sequenced on a NextSeq 550 (Illumina) with 50 bp read 1, 8 bp i7 index read, 16 bp i5 index read, and 50 bp read 2. The KO library was sequenced a second time to increase read depth, and both runs were combined during analysis.

Sample demultiplexing and initial scATAC-seq analysis including read alignment and peak calling were performed using cellranger-atac mkfastq and cellranger-atac count, respectively, using v1.2.0 software and the associated mm10 genome reference (‘refdata-cellranger-atac-mm10-1.2.0’; 10X Genomics). Count data for each sample were imported into Seurat v4.0.4 (Butler et al., 2018) in R version 4.1.1 using cell barcodes discovered by the cellranger-atac pipeline. The two datasets were merged, normalized, and clustered using Signac v1.3.0 (Stuart et al., 2020). Differential peak accessibility and gene activity scores (accessibility within gene bodies and promoter areas) were calculated for each initial cluster and were used to assign cell identities. Clusters were then merged by cell identity. Differential peaks and gene activities were re-calculated for the merged clusters. GREAT analysis was used to calculate distances of the peaks to transcription starting sites and extract affected nearby genes (McLean et al, 2010). Gene ontology and Reactome analyses were performed using g:Profiler (Raudvere et al, 2019).

## QUANTIFICATION AND STATISTICAL ANALYSIS

Quantification and statistical analyses of GRAB-HTR6, ciliary RhoA FRET/FLIM, ATAC-Airy, and scATAC-seq were described above in their respective sections. Details of experiments, including sample size and statistics can be found in the text, figures, or figure legends. Except for Figure 3G-H, all statistical tests were done using estimation statistics (Ho et al, 2019) or permutation test comparing different distributions (Figure 3H). In most cases, conventional p-values were also provided. No method was used to predetermine sample size (N), which represents data collected spanning different sessions. Blinding was not performed. Formal randomization techniques were not used.

### Quantification of cilia trajectory in the brain

3D stacks of ADCY3-stained cilia images were projected along the z axis using maximum intensity projections and analyzed with the OrientationJ plug-in in ImageJ/Fiji using a gaussian gradient (Püspöki et al., 2016; Rezakhaniha et al., 2012). The algorithm computes the structure tensor for each pixel in the image using a sliding gaussian analysis window. A local window size of 2 and 20 pixels are used for visualization and weighted histogram calculation, respectively. Gaussian fitting of the weighted histogram was performed and plotted in Graphpad Prism (v9.2).

### Quantification of cilia and serotonergic axon, synaptophysin vesicle apposition and nuclear adducin puncta

#### Linear Unmixing

3D stacks of Hoechst-stained nuclei, SERT-stained serotonergic axons (CF488), synaptophysin-stained presynaptic terminals (Alexa Fluor Plus 555), and ADCY3-stained cilia (CF633) images were acquired using FAST Airyscan. The red and far-red channel Airyscan data (41 nm x 41 nm x 189 nm voxel size, 3D Airyscan processing with Auto Filter) were unmixed via a custom MATLAB script.

#### Nucleus, cilia, axon and synaptophysin segmentation

The multichannel volumes acquired were resampled to generated isotropic voxel sizes of 189 nm. The nucleus, cilia, and axon signals were segmented using Otsu’s methods to threshold, followed by dilation and erosion operations to join fragmented structures, and finally filtered to remove small discrete objects with no biological relevance. Nuclei that were not part of the pyramidal neuron cell layer were removed by dilating the dense collection of nuclei, and subsequently retaining just the largest connected object (corresponding to pyramidal neuron cell layer). Only nuclei and associated cilia with the pyramidal neuron cell layer were included for downstream analysis steps. The central axes of segmented cilia and axon masks were computed (skeletonized), which also served to separate and identify distinct cilia. The cilia near the imaged volume boundaries or those which were associated with non-pyramidal neuron cell layer nuclei were excluded from further analyses. The synaptophysin and ADD1 puncta positions were determined by detecting their local signal maxima using a 3D Laplacian-of-Gaussian filter described previously (Aguet et al., 2016). Nucleus centers were determined by eroding the segmented nuclei masks, followed by calculating the distance transformation and thresholded to separate touching nuclei. The centroids of these discrete eroded nuclei were used to define the centers of search for ADD1 puncta with varying radii between 1 - 8 µm.

#### Calculation of axon-cilia and SERT+ synaptophysin -cilia distance

3D distance transformations of the nuclei, skeletonized axon, and synaptophysin masks were calculated and multiplied by the skeletonized cilia masks. The resulting mask encoded the distance information for the respective transformation mask in the medial axes of the cilia. As serotonergic varicosities are between 1 and 3 μm (Alvarez et al., 1998), we used 2 μm between skeletonized cilia and skeletonized axons as the cutoff for axon-contacting and non-axon-contacting cilia. Subsequently, synaptophysin puncta within 1 μm of skeletonized serotonergic axon central axes are considered associated with serotonergic axons (SERT-positive). For the visualization of the distance-encoded cilia, the cilia central axes were dilated using a sphere morphological structural kernel with a radius of 1 pixel. The resulting volumes were then projected to only visualize the distance information using a custom lookup table (Figure 3C-D). Furthermore, the total cilia length and the corresponding distance to the closest axon was plotted as a 2D density scatter using MATLAB (Chung, 2021) with a marker size set to 10 and a “jet” colormap (Figure 3F-G). Violin plots of cilium length for contacting and non-contacting cilia (Figure 3H) were generated using PlotsOfData (Postma and Goedhart, 2019). Statistical comparison of the two groups were done in GraphPad Prism. Permutation tests to compare these two distributions were performed using Mlxtend in Python (Raschka, 2018).

---

**Figure S1.**
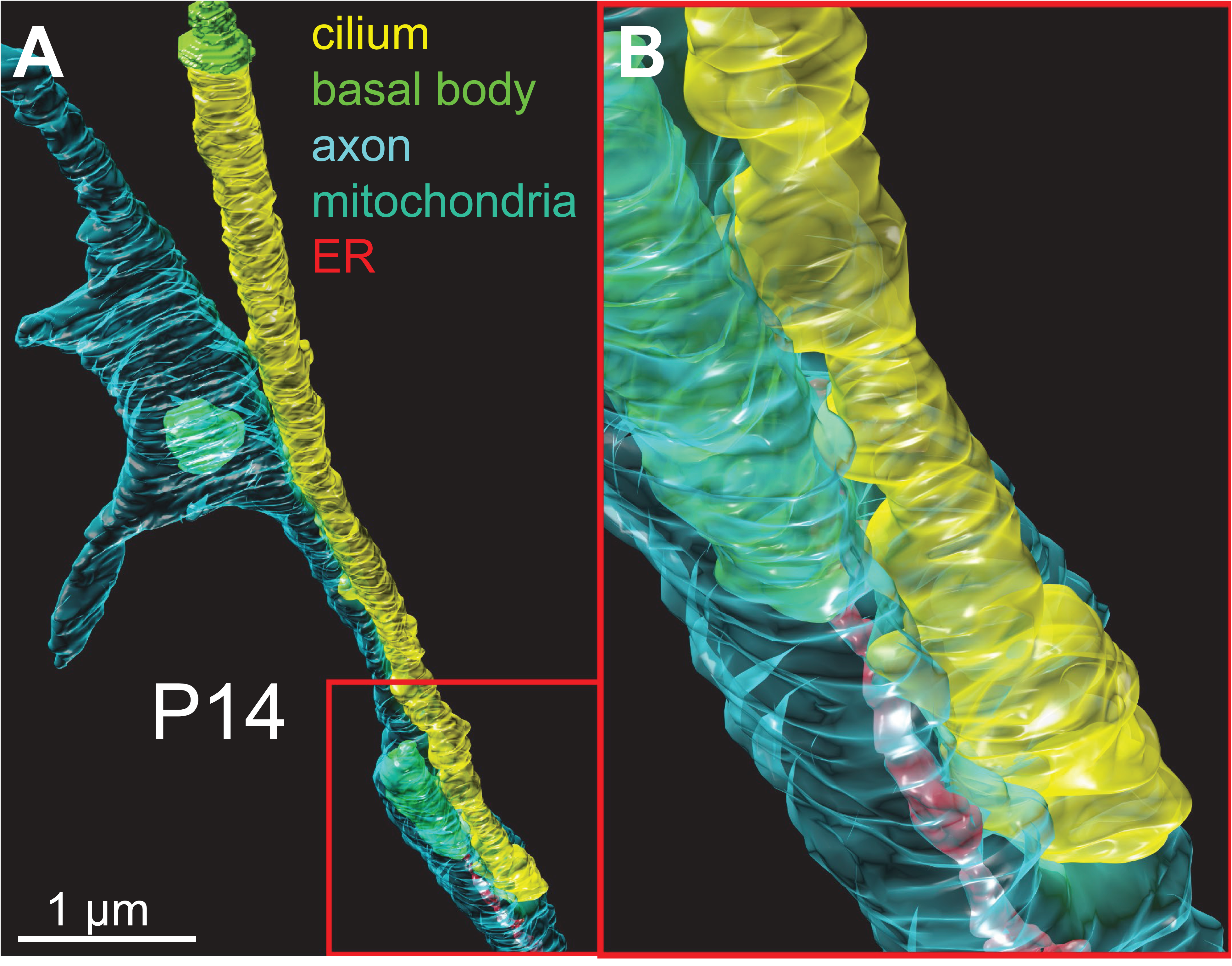
FIB-SEM reconstruction of P14 (juvenile) CA1 pyramidal neuron. Related to Figure 2. The neuronal cilium fasciculates with an axonal process. Like adult pyramidal neuronal cilia, axo-ciliary synapses are apparent. Yellow: cilium, cyan: axon, bright green: basal body, red: axonal endoplasmic reticulum, green: axonal mitochondria.

**Figure S2.**
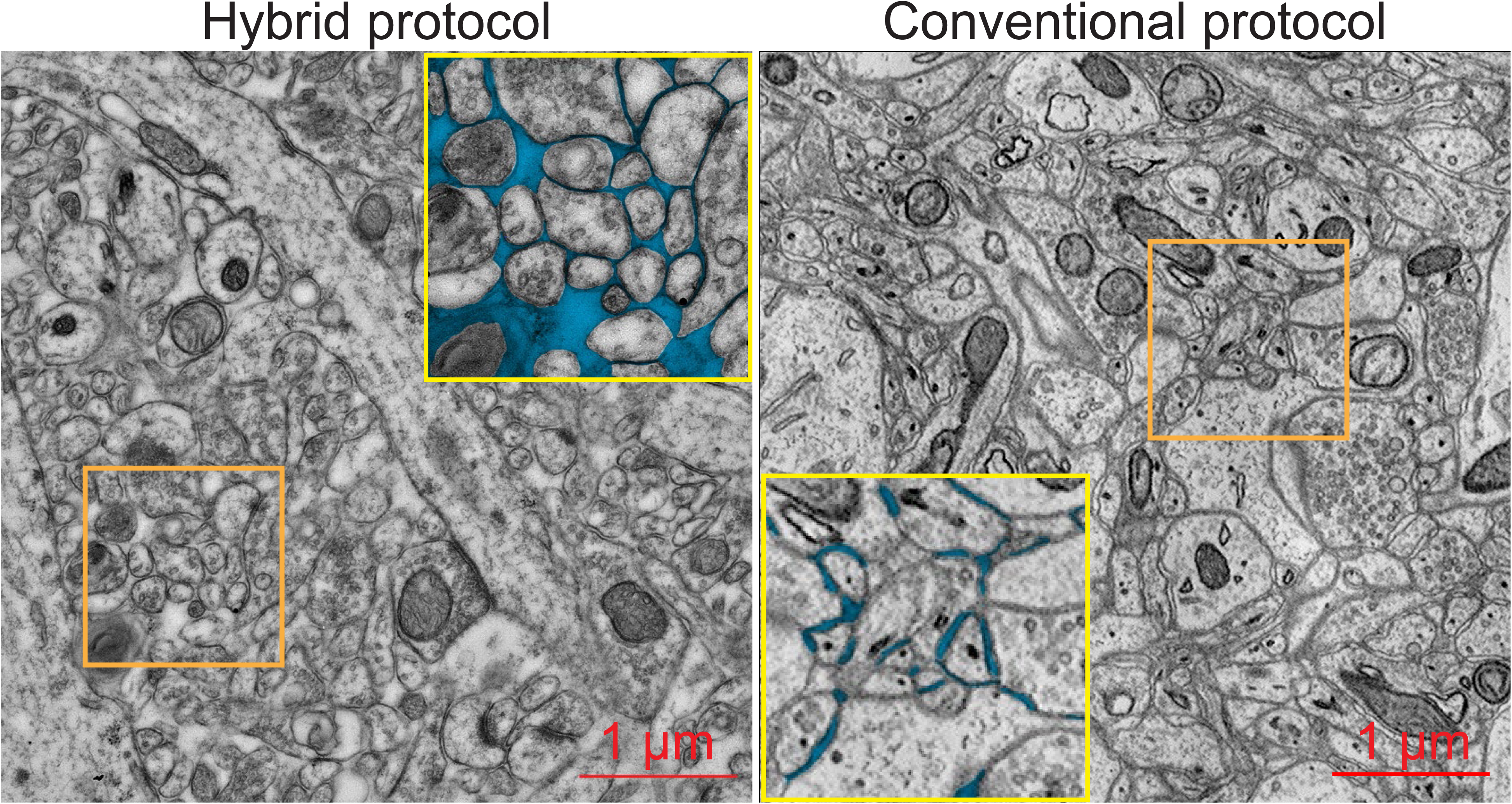
Hybrid fixation protocol preserves extracellular space. Related to Figure 2. In contrast to conventional glutaraldehyde perfusion protocols (Kasthuri et al., 2015; right, orange box magnified in the inset with yellow border), the novel hybrid protocol preserves the extracellular space (orange box magnified in the inset with yellow border). Note the rounded morphology of neuronal processes and significantly greater amount of extracellular space (blue). See Methods for details.

**Figure S3.**
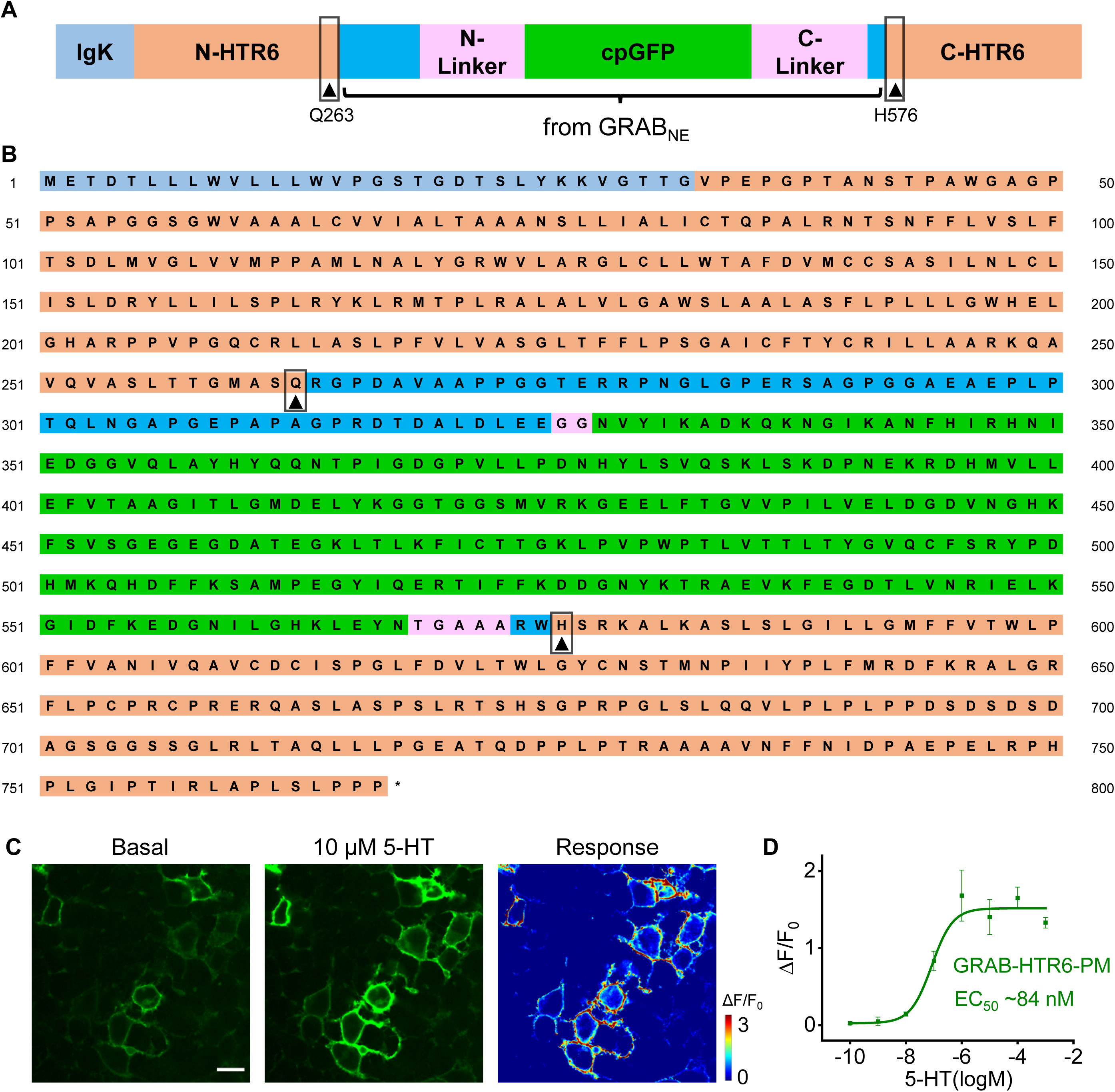
The amino acid sequence and characterization of the GRAB-HTR6-PM sensor. Related to Figure 4. (**A**) Schematic diagram of GRAB-HTR6-PM sensor’s structure. The cpEGFP and linkers were transplanted from GRAB_NE_. (**B**) The amino acid sequence of the GRAB-HTR6-PM sensor. The numbering used in the figure starts from the IgK leader sequence. Insertion sites Q263^ICL3^ and H576^6.26^ are indicated by black arrowheads below relevant amino acids. (**C**) Representative images show the expression of GRAB-HTR6-PM sensor (left, before application of 10 μM 5-HT, middle, after application of 10 μM 5-HT) and the response (right, pseudocolor) in HEK293T cells. Scale bar, 20 μm. (**D**) The dose-response curve of the HTR6-GRAB-PM sensor tested in HEK293T cells; EC_50_ labeled. Data are shown as mean ± S.E.M. and n = 3 wells with 300‒500 cells per well.

**Figure S4.**
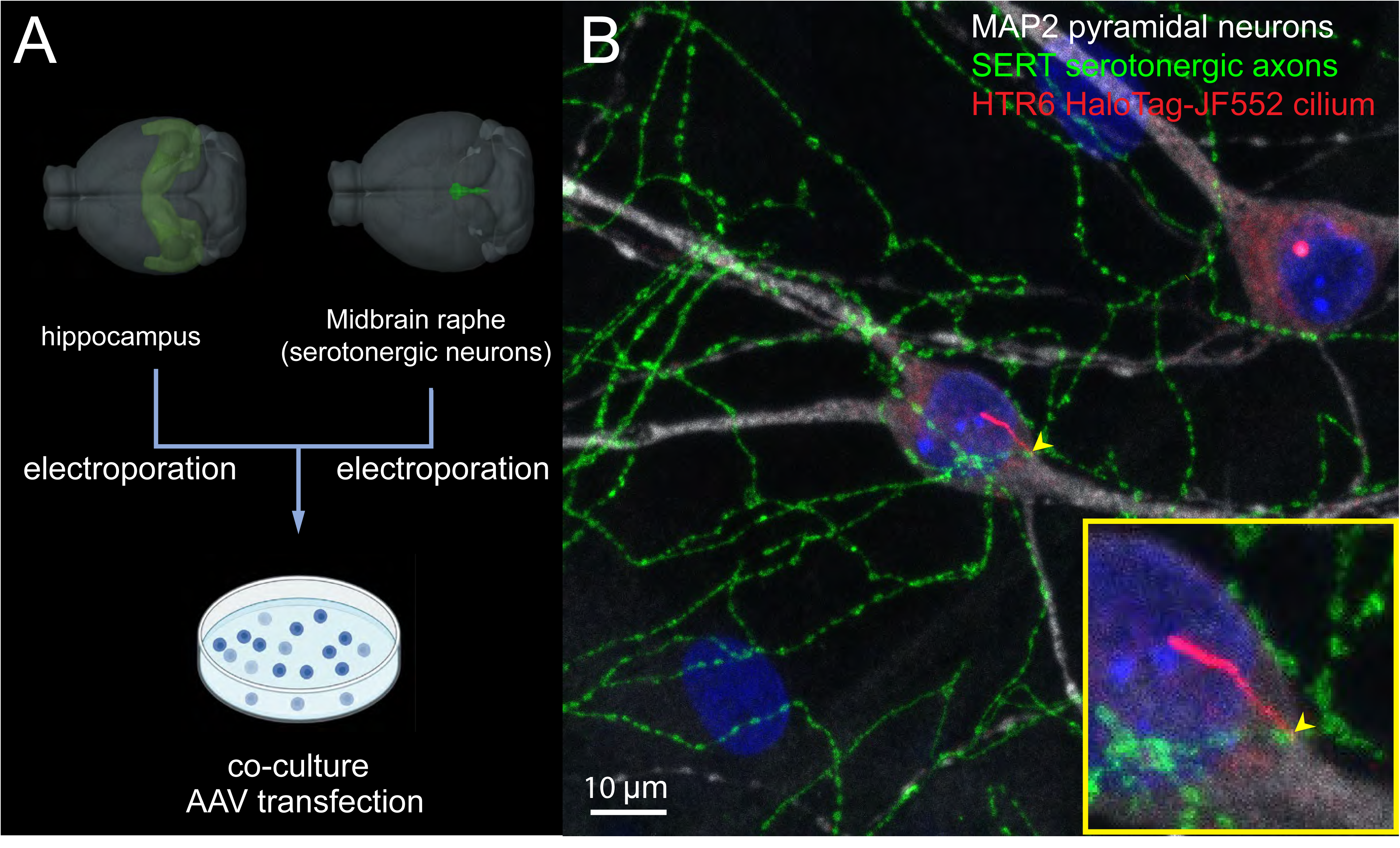
Serotonergic axo-ciliary synapse *in vitro*. Related to Figure 4. (**A**) Overall workflow. Hippocampal and raphe neurons were dissociated from the hippocampus and midbrain, respectively. In some experiments, constructs were electroporated separately before co-plating in the same well (see Methods). (**B**) Serotonergic axo-ciliary synapses occur in vitro. On average (1 million cell total, 300,000/cm^2^ density, 1:1 hippocampal and midbrain cell ratio), there are 5-10 serotonergic neurons and ∼5 axo-ciliary synapses per well.

**Figure S5.**
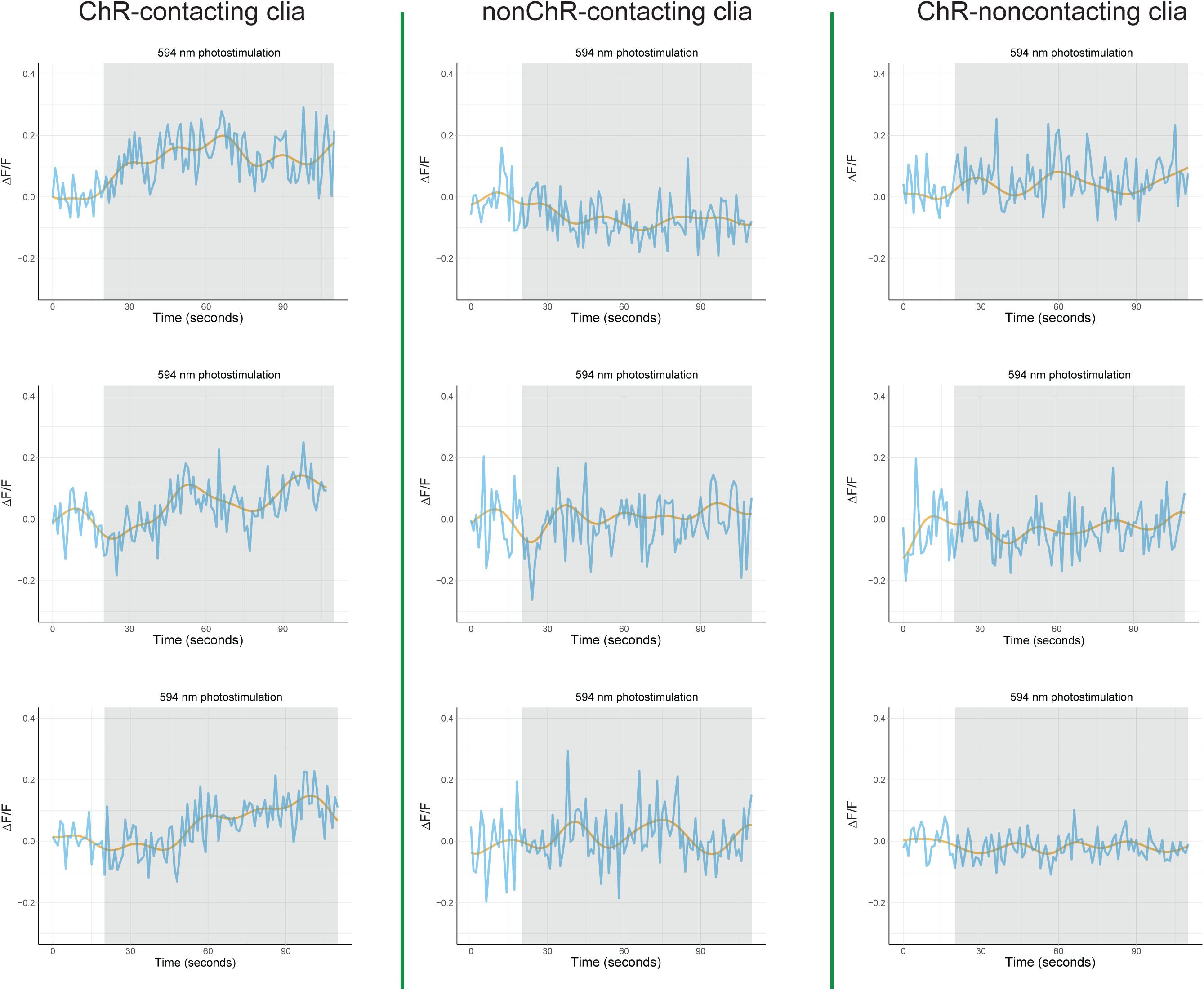
Measurement of GRAB-HTR6-cilia of axo-ciliary synapses. Related to Figure 4. Fluorescence intensity changes during optogenetic stimulation (starting at 20 s) across three different groups. Blue: raw trace, orange: raw trace filtered by a low pass filter (0.2 Hz). The peaks of the filtered traces during photostimulation were used for statistical testing shown in Figure 4C.

**Figure S6.**
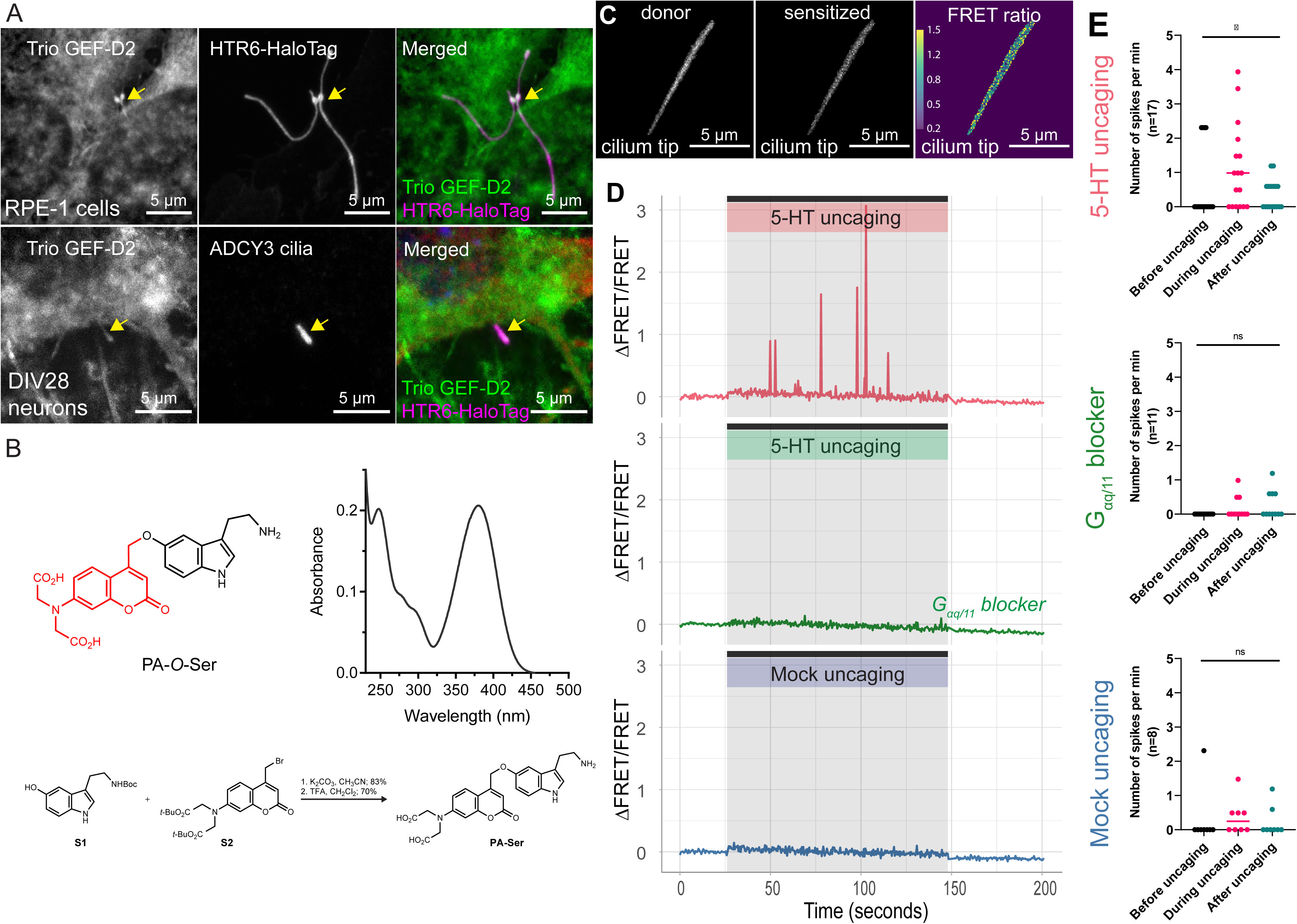
Ciliary G_αq_-Trio-RhoA signaling in RPE-1 cells. Related to Figure 5. (**A**)Trio is present in HTR6-cilia in RPE-1 cells and WT cultured hippocampal neuronal cilia. Top panel: RPE-1 cells stably expressing the Tet-inducible HTR6-HaloTag. HaloTag was labeled with Janelia Fluor 552 (magenta in the merged panel), fixed, and stained with an antibody against the Trio GEF-D2 domain (green, merged panel). Lower panel: Trio is present in WT cultured hippocampal neuronal cilia. DIV28 cultured rat hippocampal neurons were fixed and immunostained with anti-ADCY3 antibody (neuronal cilia marker, magenta; merged panel), anti-MAP2 antibody (neuronal marker, red; merged panel), Hoechst 33342 (nucleus, blue; merged panel), and anti-Trio GEF-D2 antibody (green, merged panel). The Trio GEF-D2 signal in both cases was amplified with the Alexa 488 tyramide signal amplification system (Thermo Fisher Scientific). Images were processed with the Subtract Background (50 pixels with sliding paraboloid) algorithm in ImageJ/Fiji to enhance contrast for qualitative demonstrations. (**B**) Properties of photo-activatable (“caged”) serotonin (PA-Ser). Top left: Chemical structure of PA-Ser. Top right: Absolute absorption spectrum of a solution of PA-Ser (10 μM) in PBS. This molecule displayed an absorption maximum of 380 nm with an extinction coefficient (ε) of 21,100 M^−1^cm^−1^; the relatively broad absorption spectrum gives substantial absorption at 405 nm (ε = 12,100 M^−1^ cm^−1^). Upon photolysis, PA-Ser releases ∼10% of serotonin along with other major photoproducts generated primarily via a photo-Claisen pathway (Wong et al., 2017). Lower panel: Synthesis of PA-Ser through alkylation of Boc-protected serotonin (S1) with {7-[bis(carboxymethyl)amino] coumarin-4-yl} methyl (BCMACM) bromide (S2). This coumarin-based BCMACM photolabile group exhibits high aqueous solubility and relatively large one- and two-photon action cross-sections (Hagen et al., 2008). (**C-E**) Serotonin stimulation of ciliary HTR6 activates RhoA. RPE-1 cells stably expressing a Tet-inducible HTR6-RhoA FRET based sensor. **C**. Donor emission (sGFP2), sensitized emission (mScarlet-I), and FRET ratio calculated by dividing sensitized emission by donor emission of a single cilium. Local serotonin uncaging at 0.5 Hz results in RhoA activity spikes (**D** Top panel shows a sample trace, quantified in **E** top panel, p value=0.04). The effect is largely attenuated by pre-treating samples with the Gαq/11 blocker, YM-254890 (1 μM, **D** middle panel is a sample trace, quantified in **E** middle panel, p value=0.12). Mock uncaging had minimal effect on the RhoA FRET ratio (**D** lower panel is a sample trace, quantified in **E** lower panel, p value =0.52). For **E**, the spikes are defined as ΔF/F greater than or equal to 0.52 (the mean ± 3 S.D. in the 5-HT uncaging measurements). Horizontal lines represent the median values. Statistical tests comparing before, during, and after uncaging used the Friedman test (non-parametric).

**Figure S7.**
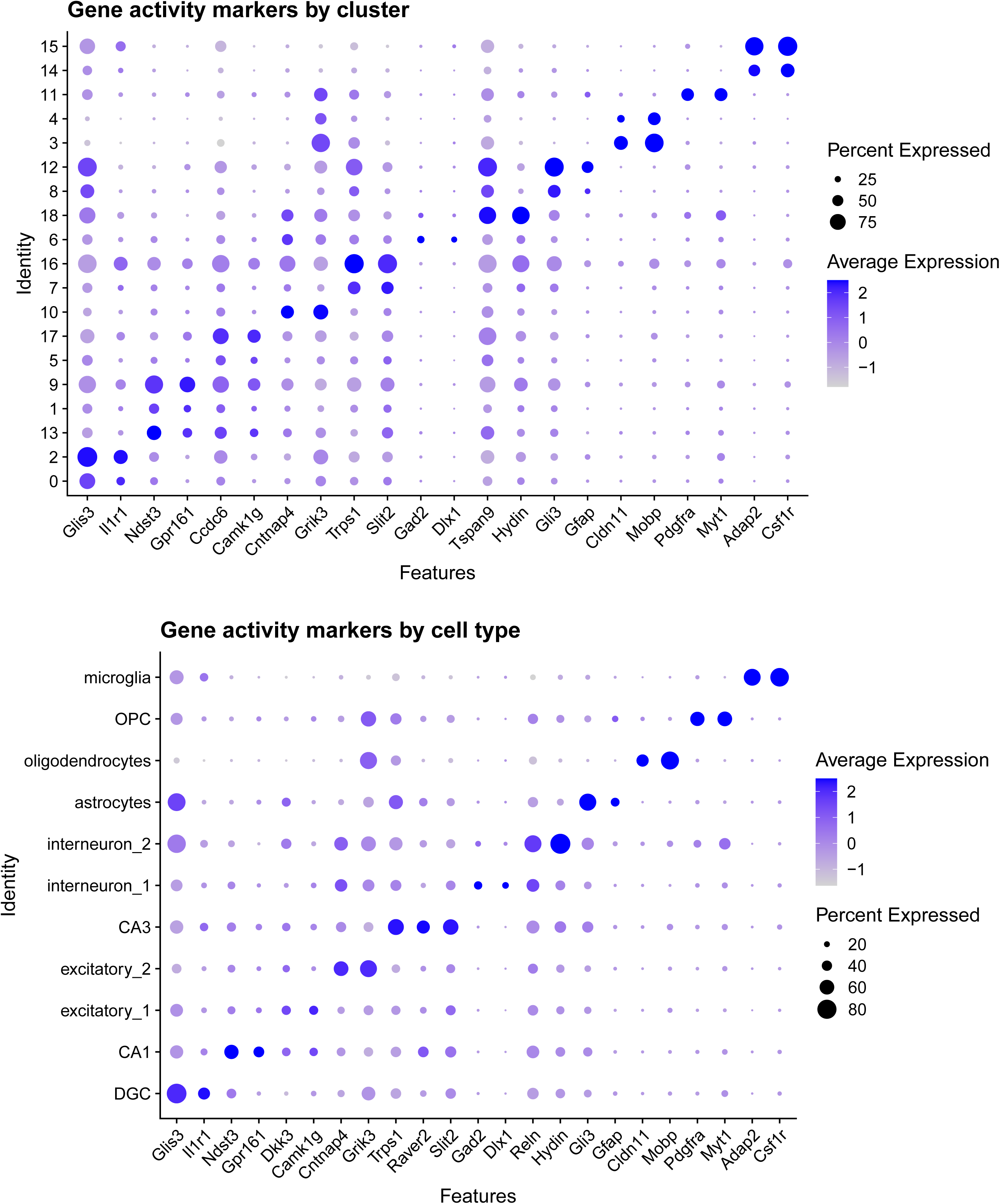
Marker genes for cell type assignment. Related to Figure 7. For each individual marker gene, we plotted the average gene activity level (see Methods) and percent of cells that expresses the gene. Top panel: original clusters; bottom panel: merged clusters.

## Graphical Abstract

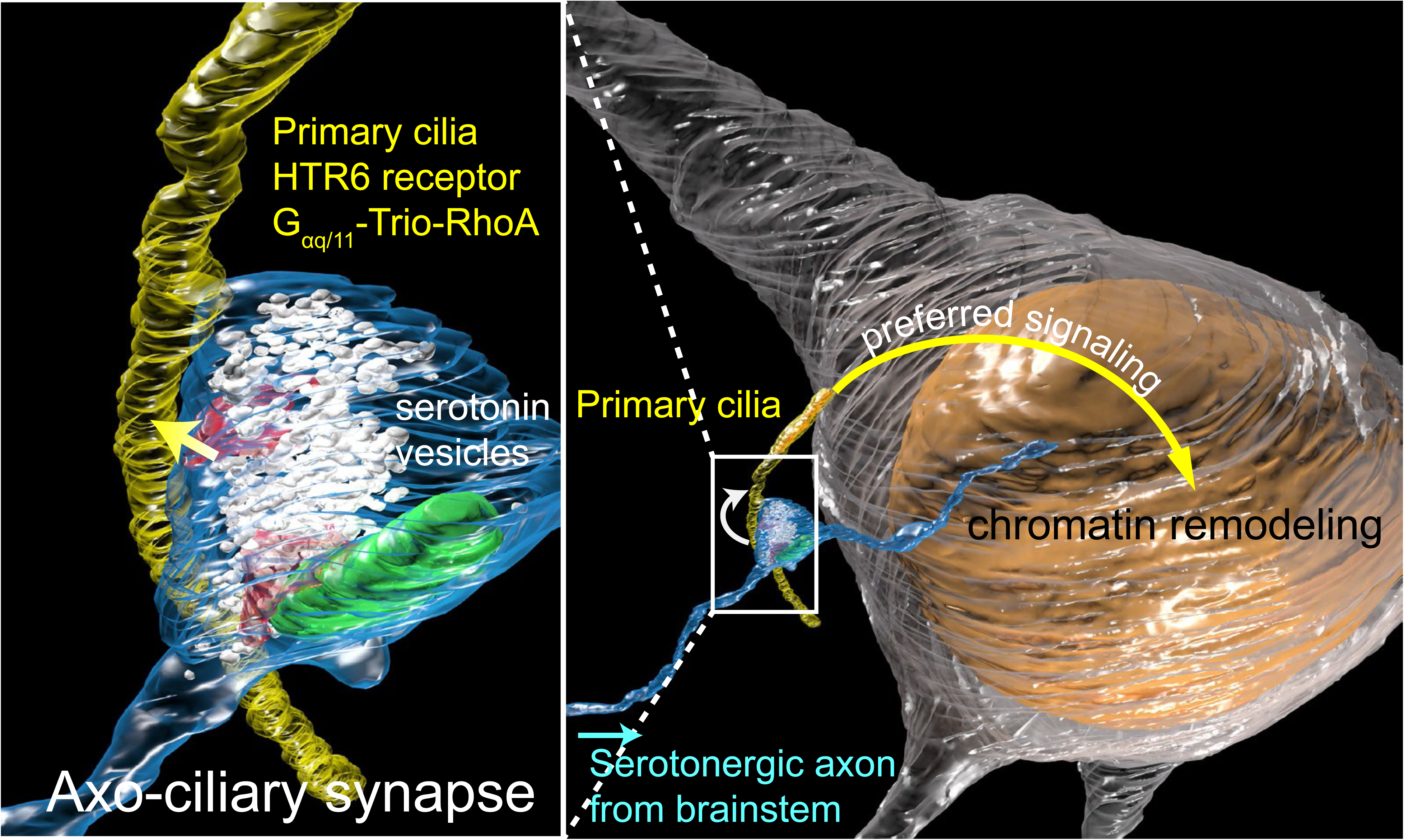
Model of the serotonergic axo-ciliary synapse. Identification of synapses between serotonergic axons and pyramidal neuron primary cilia. Cilia-specific serotonin receptor constitutes a preferred signaling pathway to the nucleus. Loss of this pathway results in chromatin remodeling.

